# Regulation of plant metabolism under elevated CO_2_

**DOI:** 10.1101/2024.08.23.609313

**Authors:** Danial Shokouhi, Jakob Sebastian Hernandez, Dirk Walther, Gabriele Kepp, Serena Schwenkert, Dario Leister, Jürgen Gremmels, Ellen Zuther, Jessica Alpers, Thomas Nägele, Arnd G. Heyer

## Abstract

Plant responses to changing environments afford complex regulation at transcriptome and proteome level to maintain metabolic homeostasis. Homeostasis itself constitutes a complex and dynamic equilibrium of metabolic reactions and transport processes among cellular compartments. In the present study, we aimed at the highest possible resolution of this network by combining analysis of transcriptome, proteome and subcellular resolved metabolome of plants exposed to rising carbon dioxide concentrations over a time course of one week. To prove suitability of our approach, we included mutants affected in photorespiratory metabolism and, thus, should deviate from the wildtype in their response to elevated CO_2_. Our multi-omics analysis revealed that the *hpr1-1* mutant, defective in peroxisomal hydroxypyruvate reduction, is also affected in cytosolic pyruvate metabolism, reaching out to cysteine synthesis, while the hexokinase mutant *hxk1* displays a disturbed redox balance upon changing CO_2_ levels. For the third mutant, defective in the mitochondrial protein BOU, we found compelling evidence that the function of this transporter is related to lipoic acid metabolism, thus challenging current interpretations. This demonstrates that the combined omics approach introduced here opens new insights into complex metabolic interaction of pathways shared among different cellular compartments.

## Introduction

Studying regulation of plant metabolism under changing atmospheric conditions is complicated by the high level of compartmentalization of plant cells, but also by the redundancy of pathways in different compartments, e.g., glycolysis, oxidative pentose phosphate pathway or synthesis of various amino acids in the cytosol, plastids and mitochondria. To disentangle overlapping reactions, mutants lacking specific isoforms of enzymes are frequently utilized that disrupt metabolite conversion in a specific compartment. While this strategy enabled insight into the complex interactions of metabolic pathways, knowledge is still limited due to high intracellular mobility of metabolic intermediates and also because of so-called pleiotropic effects of mutations, which are unexpected based on current knowledge of functions of the mutated genes ^*1*^.

In the present study we aimed at combining data sets from the transcriptional and proteomic level with metabolite profiles for different subcellular compartments in a time series on plants exposed to an increased CO_2_ concentration. By applying the highest analytical resolution possible by now, we asked whether this could solve the problem of pleiotropy. We used three different mutants of Arabidopsis reported to be affected in photorespiratory metabolism.

Photorespiration is, next to photosynthetic CO_2_ fixation within the Calvin-Benson-Bassham cycle (CBBC), the second most important metabolic route in green tissues of plants with regard to turnover during the light phase and has an important function also in nitrogen acquisition ^*2*^. It involves at least four different cellular compartments, i.e. plastids, cytosol, peroxisomes and mitochondria, but recent findings point to a function of the vacuole in photorespiration, too ^*3*^.

Photorespiration is initialized by oxygenation of ribulose-1,5-bisphosphate that results from the dual use of substrates, carbon dioxide and oxygen, by the enzyme Ribulose-Bisphospate-Carboxylase/Oxygenase (Rubisco). Oxygenation leads to the production of phosphoglycolate, which inhibits activity of enzymes in the CBBC ^*4*^. After dephosphorylation, glycolate is oxidized to glyoxylate and transaminated to yield glycine in the peroxisomes. Two molecules of glycine are converted to serine in the mitochondria, releasing carbon dioxide and ammonia, which must be re-fixed into glutamine in the plastids. Serine is deaminated to hydroxypyruvate again in the peroxisome, which is then reduced to glycerate that, after phosphorylation, can re-enter the CBBC in the plastids. The *hpr1-1* mutant of Arabidopsis is impaired in the last step of the pathway, i.e. reduction of hydrxypyruvate, and is one of the few examples of photorespiratory mutants able to survive at ambient CO_2_ concentration, at least in part due to the presence of a cytosolic isoform of the enzyme ^*5*^. While HPR is directly involved in the photorespiratory cycle, hexokinase-1 (HXK1) is involved in the phosphorylation of glucose and fructose produced from sucrose by the action of invertase and has a role as a sugar sensor in plant metabolism ^*6, 7*^. Its involvement in photorespiration has been concluded from experiments with the *hxk1* mutant under high light intensity, where increased photorespiratory turnover led to an accumulation of serine in mitochondria of wild type plants, while it resided in the cytosol of *hxk1* ^*8*^. The third mutation in the set is “a bout de souffle” (*bou*, out of breath), which is blocked in development under ambient CO_2_ concentration, thus showing a severe photorespiratory phenotype, accompanied by a very high glycine-to-serine ratio ^*9*^. The BOU enzyme is a mitochondrial transporter the substrate of which is not unequivocally clear. *In vitro* evidence for transport of glutamate has been presented, and it has been hypothesized that its knockout might interfere with poly-glutamylation of dihydrofolate reductase that is needed for glycine decarboxylation ^*10*^. However, the mutant shows a very complex phenotype, and the presence of additional mitochondrial glutamate transporters ^*11*^ might argue against a lack of glutamate as sole reason for the mutant phenotype.

In the present study, we aimed at analyzing changes in metabolic regulation resulting from increased atmospheric CO_2_ concentration. To this end, we transferred plants from ambient (450 ppm) to about two-fold higher (1000 ppm) CO_2_ levels and recorded changes in gene expression, protein abundance and compartment-specific metabolite levels over a time course of seven days. This experimental design allowed for an investigation of the effects of environmental changes related to the anthropogenic climate change. In addition, organelle interaction that was affected by the change in the ratio of photosynthesis to photorespiration could be analyzed. Combined transcriptomics, proteomics, and compartment specific metabolomics provided detailed information on mutant effects and enabled new understanding of metabolic interactions in a changing environment.

## Results and Discussion

### The metabolic phenotype of *hpr1-1* mutants comprises strong effects on cytosolic glycerate and pyruvate metabolism

To gain a detailed understanding of how the metabolic mutants were affected in primary metabolites with relation to the photorespiratory pathway, the *OmicsDB* was created in conjunction with an application programming interface to allow for fast data exploration and straightforward hypothesis testing (see Methods). Extracting subcellular metabolome information with direct relation to photorespiration revealed that *hpr1-1* plants showed the expected strong photorespiratory metabolic phenotype under ambient CO_2_ (aCO_2_) which clearly separated them in a principal component analysis (PCA) from all other genotypes (Supplementary Figure S1). After transfer to elevated CO_2_ (eCO_2_), this effect was diminished but *hpr1-1* samples still clearly separated from Col-0. Both, glycine and serine had a high influence on separating *hpr1-1* from Col-0 under aCO_2_ (Supplementary Figure S1 B). However, to a similar extent, elevated glycerate levels, especially in mitochondria, differentiated *hpr1-1* from all other genotypes (Figure 1). This was counter-intuitive, because HPR1 contributes to glycerate biosynthesis, and thus a reduction in this metabolite could be expected. A similar, yet unexplained, observation was already made at whole-cell level and was shown not to depend on the cytosolic isoenzyme HPR2 ^*5*^.

**Figure 1.**
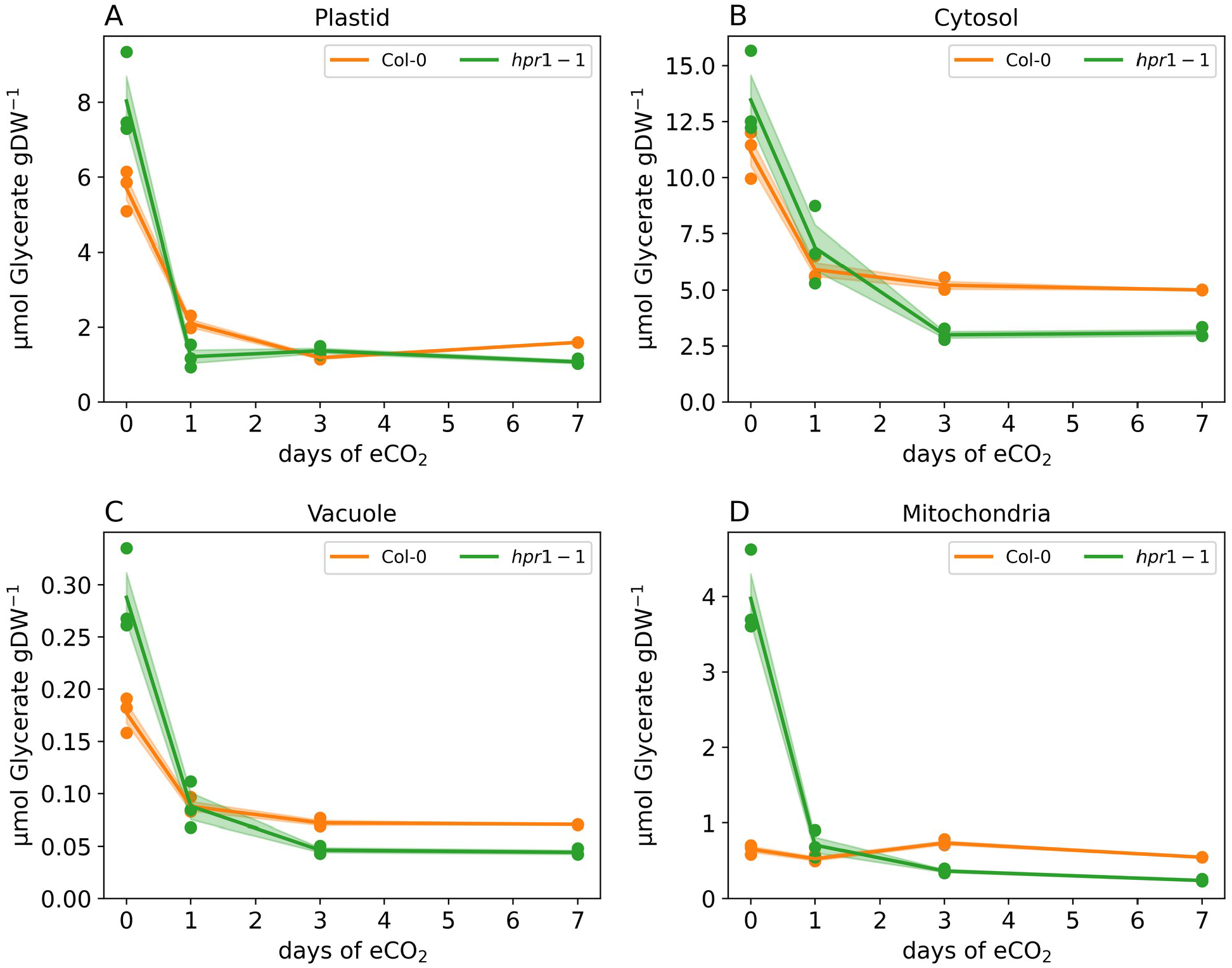
Compartment specific glycerate dynamics. Ordinates reflect glyceric acid amounts in µmol gDW^-1^, abscissae show time of exposure to eCO_2_ in days. (A) plastid; (B) cytosol; (C) vacuole, (D) mitochondria. Genotypes are indicated by color (orange – Col-0; green – *hpr1-1*). Filled circles indicate mean values, shaded areas indicate standard deviation, (n = 3)

Compartment-specific metabolomics revealed low glycolate in plastids, but accumulation in the cytosol of *hpr1-1* at aCO_2_ (Supplementary Figure 2 A). This suggested an alternative, so-far undescribed cytosolic route in *hpr1-1* from glycolate to glycerate, which then could be transported into the plastid, probably via the plastidial glycolate /glycerate antiporter PLGG1 (AT1G32080) or the bile acid/sodium symporter BASS6 (AT4G22840). Compared to Col-0, transcripts of PLGG1 were slightly elevated in *hpr1-1* and immediately responded to eCO_2_, suggesting a direct photorespiratory role. The alternative pathway would involve cytosolic pyruvate and cysteine, both being strongly elevated in *hpr1-1* at eCO_2_ (Figure 2 A, B). Cysteine dynamics were uncoupled from serine and glycine (Figure 3 A), indicative of an alternative metabolic role of cysteine as an intermediate in the pathway from serine to glycerate, thus bypassing hydroxypyruvate (Supplementary Figures S3, S4). Key enzymes on this route, O-acetylserine-thiolyase (OASA1) and L-cysteine desulfhydrase 1 (DES1 - AT5G28030), which catalyzes the interconversion of cysteine to pyruvate releasing ammonia and sulfide, were upregulated at transcript level (log2FC: 0.532; adj. p val. < 0.00001 for one sided z-test, n = 3; Figure 3 B, C).

**Figure 2.**
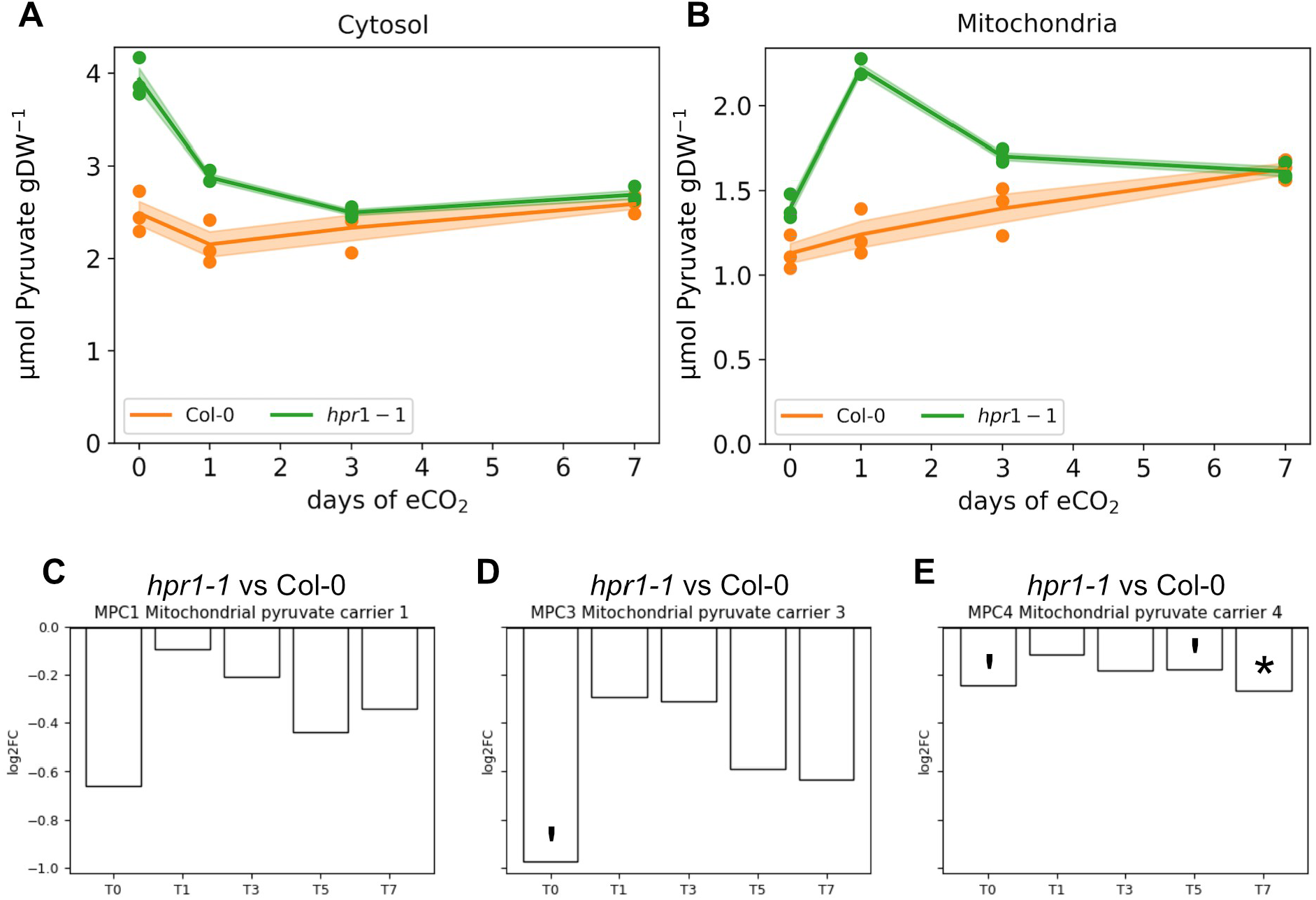
Subcellular pyruvate concentration of the cytosol and mitochondria, with log2FCs of associated transporters. **(A)** Dynamics of cytosolic pyruvate concentration, **(B)** Dynamics of mitochondrial pyruvate concentration. Ordinates reflect pyruvate amounts in µmol gDW^-1^ (mean ± SD; n = 3), abscissae show time of exposure to eCO_2_ in days. **(C) – (E)** Log2FC between *hpr1-1* and Col-0 for MPC1, MPC3 and MPC4, respectively (* - p value < 0.05, ‘ - p value < 0.1; n = 3).

**Figure 3.**
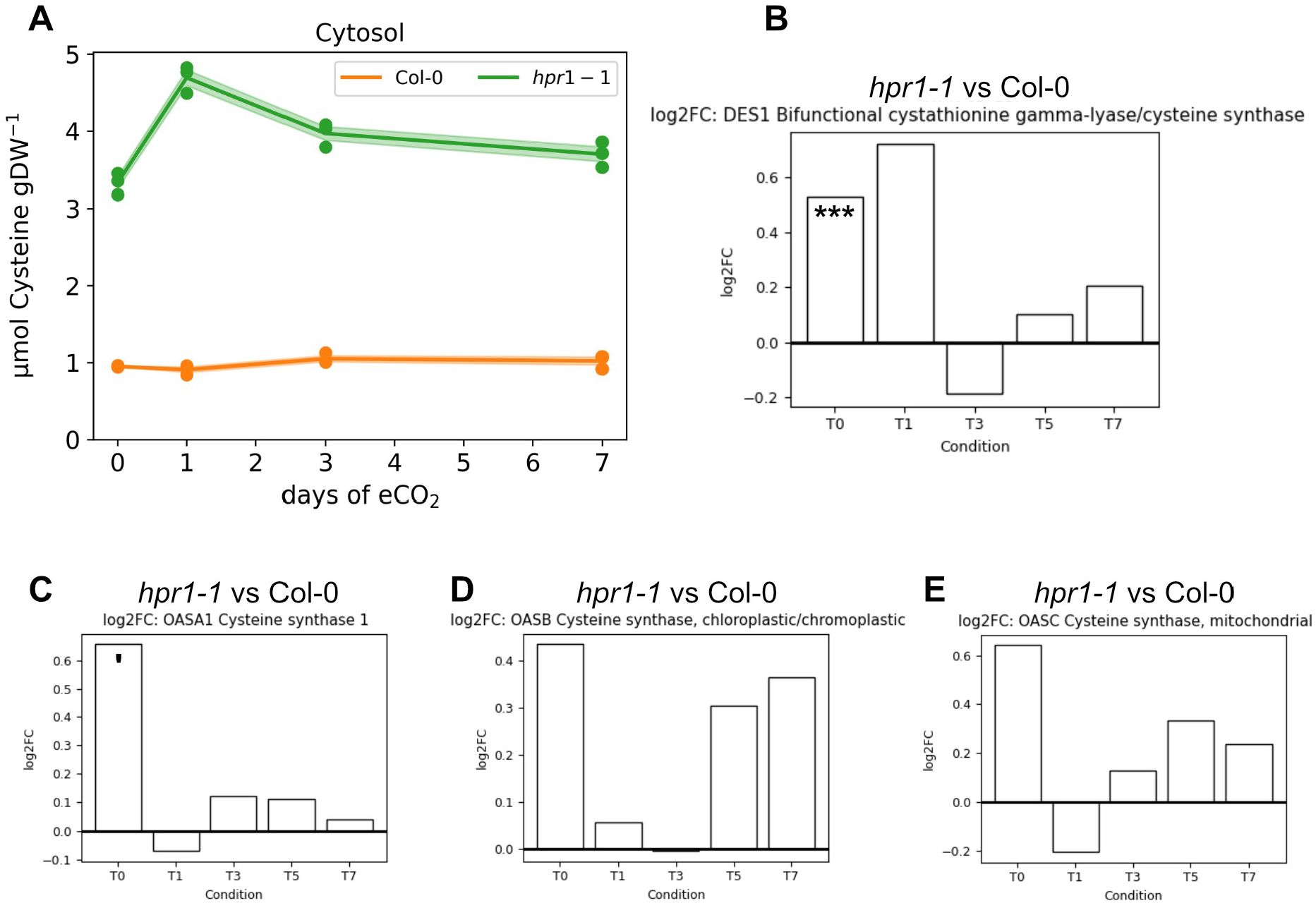
Cytosolic cysteine dynamics with associated enzymes. **(A)** Dynamics of cytosolic cysteine concentrations. Ordinate reflects cysteine amounts in µmol gDW^-1^ (mean ± SD; n = 3), abscissa shows time of exposure to eCO_2_ in days. **(B)** Log2FC of DES1 comparing *hpr1-1* to Col-0, **(C) – (E)** Log2FC of enzymes for cysteine biosynthesis enzymes. replicates (*** - p value < 0.001, ‘ - p value < 0.1; n = 3).

Cytosolic pyruvate, a product of glycolysis, is supposed to be transported into mitochondria as substrate for the TCA cycle. If involved in the aforementioned pathway, reduced influx into mitochondria would be expected. Indeed, mitochondrial pyruvate carriers 1, 3 and 4 (MPC1 - AT5G20090, MPC3 - AT4G05590, MPC4 - AT4G22310; Figure 2 C - E) displayed slightly lower transcript levels in *hpr1-1* compared to Col-0. This trend was most pronounced in MPC3 that showed a log2FC of -0.977 (adj. p val. 0.067 for one sided z-test, n = 3) under ambient conditions. Transfer to eCO_2_ resulted in a log2FC of -0.292 after one day of exposure. In line with this, the transcript of pyruvate dehydrogenase kinase (PDK - AT3G06483), an inhibitor of the pyruvate dehydrogenase complex, was significantly upregulated (Supplementary Figure S5).

In addition to serine, malate from various sources could serve to replenish glycerate, either via pyruvate, depending on malic enzyme, or via oxaloacetate and phosphoenolpyruvate (PEP), involving malate dehydrogenase and phosphoenolpyruvate carboxykinase (PCK), thus explaining the accumulation of cytosolic malate in *hpr1-1*. Depending on the source of pyruvate and acetyl-CoA such a bypass could achieve similar carbon recuperation rates as the base photorespiratory pathway. In such a scenario, acetyl-CoA could originate from either citrate via activity of ATP-citrate lyase (ACL), or acetate depending on the compartment of cysteine synthesis. Indeed, cytosolic citrate was found to accumulate in *hpr1-*1 and transcripts for enzymes related to acetyl-CoA synthesis were found to be upregulated, too (Supplementary Figure S6). In summary, the combined Omics analysis opened the perspective of a new route to glycerate regeneration under photorespiratory conditions, which could make a substantial contribution in the *hpr1-1* mutant.

### Mutation of HXK1 affects cellular redox balance under photorespiratory conditions

An outstanding feature characterizing metabolic changes in the *hxk1* mutant at eCO_2_ was a significant increase in soluble sugars. Hexoses increased specifically in the cytosol of *hxk1* while sucrose accumulated in the vacuole, indicating an expectable slow-down in sucrose cycling as a consequence of reduced HXK1 activity.

Looking deeper into the subcellular metabolomes of Col-0 and *hxk1* indicated opposite trends of pyruvate allocation under eCO_2_ between chloroplasts and mitochondria (Figure 4). Under aCO_2_, plastidial pyruvate in *hxk1* exceeded that in Col-0, indicating different activity of plastidial glycolysis. Glycolytic and TCA cycle activity have earlier been shown to be inversely related to ambient CO_2_/O_2_ ratio and dependent on redox power ^*12*^. Although glycolytic flux in illuminated leaves may only partially depend on cytosolic hexose phosphorylation, because sugar phosphates are directly available from the triose phosphate export from plastids ^*13*^, the shift to plastidial pyruvate points out that, under aCO_2_, deficiency of HXK1 may indeed impair glycolytic flux, which would in turn affect the redox status of the NAD pool. Thus, protein levels of the gene ontology (GO) terms “cell redox homeostasis” and “TCA cycle” were correlated genotype-wise in a PCA (Supplementary Figure S7). The proteomes of all genotypes but *hxk1* showed a clear separation of aCO_2_ (time point T0) and the first day at eCO_2_ (T1), while response was substantially delayed in *hxk1*. A strong effect of the HXK1 mutation was observed for peroxisomal catalase 2 (CAT2, AT4G35090), which dissipated hydrogen peroxide produced in photorespiration. CAT2 significantly decreased within the first day of eCO_2_ in all genotypes except *hxk1*, where it declined with a significant delay until day 5 of eCO_2_ (Supplementary Figure S8).

**Figure 4.**
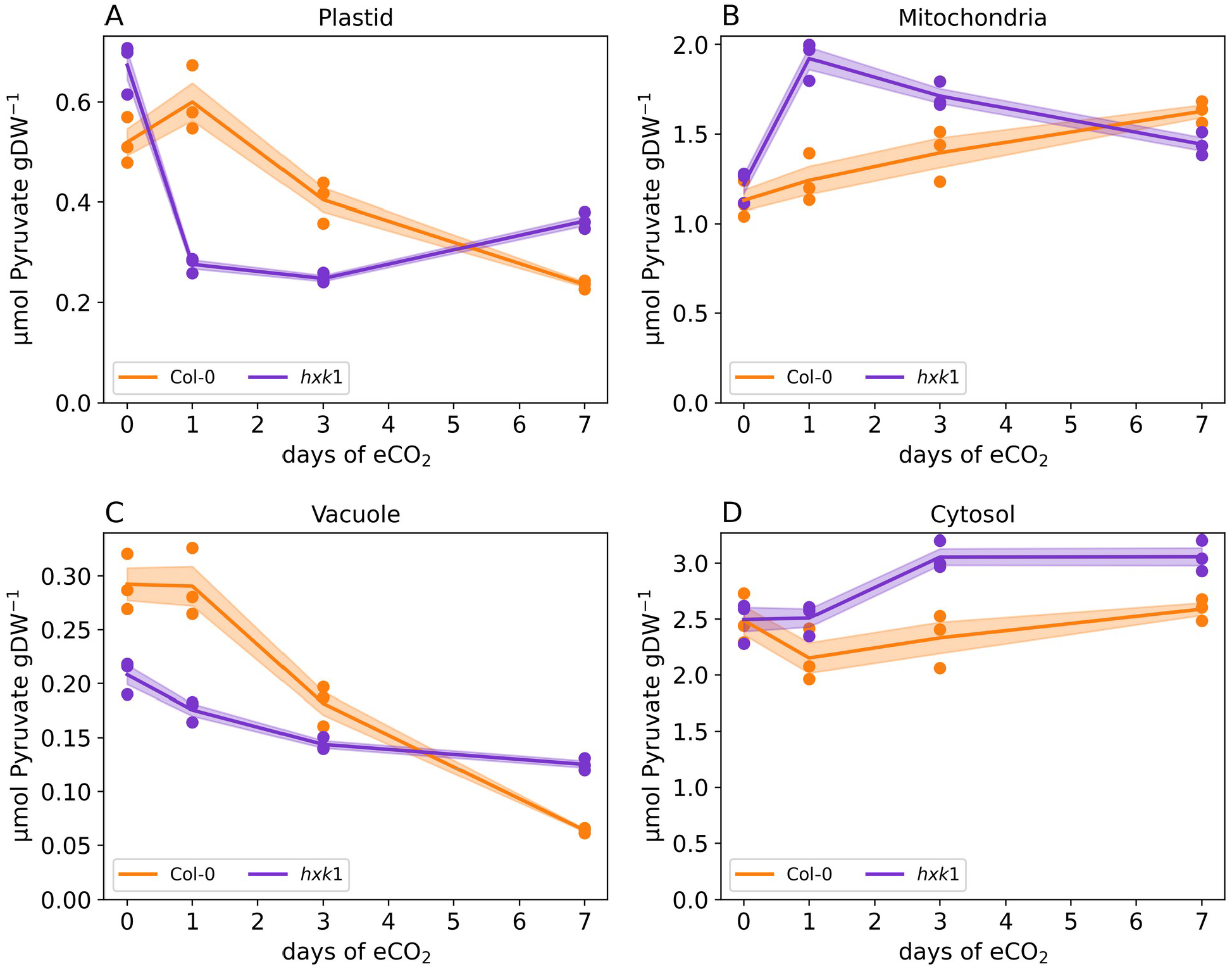
Compartment-specific pyruvate dynamics. Ordinates reflect pyruvate amounts in µmol gDW^-1^, abscissae show time of exposure to eCO_2_ in days. (A) plastid; (B) mitochondria; (C) vacuole, (D) cytosol. Genotypes are indicated by color (orange – Col-0; purple – *hxk1*). Filled circles indicate mean values, shaded areas indicate standard deviation, (n = 3).

It has been reported that, during the photorespiratory cycle, redox equivalents are shuttled between mitochondria, cytosol and peroxisomes ^*14*,^ where they are used for the reduction of hydroxypyruvate to glycerate ^*15*.^ Peroxisomal malate dehydrogenase, pMDH2 (AT5G09660), that is involved in this shuttle was down regulated in *hxk1* at early eCO_2_, but not in the wildtype (Figure 5 D).

**Figure 5.**
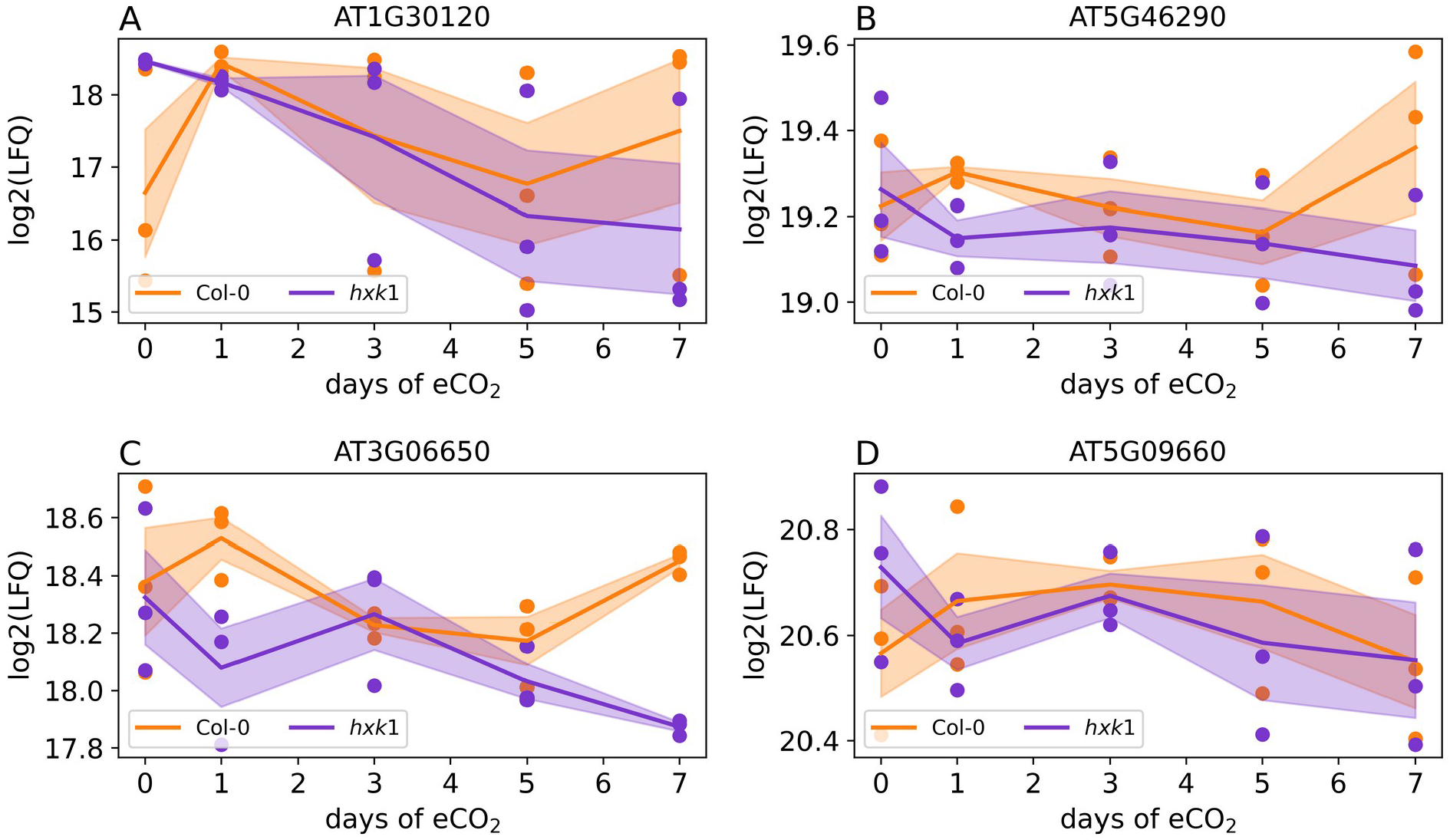
CO_2_-dependent protein dynamics of fatty acid biosynthesis and peroxisomal malate dehydrogenase. (A) plastidial pyruvate dehydrogenase E1 component (AT1G30120), (B) 3-oxoacyl-[acyl-carrier-protein] synthase I (AT5G46290), (C) ATP-citrate synthase B-1 (AT3G06650), (D) peroxisomal NAD-Malate dehydrogenase 2 (AT5G09660). orange: Col-0; purple: *hxk1*. lines and shaded areas represent means +/-SE (n = 3).

Downregulation at eCO_2_ could follow lowered photorespiratory activity, but the delayed downregulation of peroxisomal catalase CAT2 in *hxk1* during eCO_2_ exposure opens an alternative view. During photorespiration, glycolate is oxidized to yield glyoxylate thereby producing H_2_O_2_, the substrate for catalase-driven disproportionation. If hydroxypyruvate is not reduced by HPR and malate dependent pMDH2, it can non-enzymatically be oxidized to glycolate and subsequently to formate with H_2_O_2_ as oxidant ^*15*^. This reaction is relevant in the *hpr1-1* mutant, but may also take place in *hxk1*. Strikingly, both mutants show very high levels of malate in the vacuole, where it is not available as a redox shuttle to the peroxisome. This holds true for *hpr1-1* under all conditions, but only at eCO_2_ in *hxk1*.

It has been reported earlier that oxylipins interact and modulate proteins of ROS synthesis and scavenging, among others also CAT2 (an overview is provided in ^*16*^). Oxylipins are bioactive lipid derivatives which are synthesized from polyunsaturated fatty acids (PUFAs). They are involved in diverse plant-environment interactions, their functions ranging from phytohormone biosynthesis to retrograde signaling ^*16*^. While, to the best of our knowledge, until today no direct regulatory interaction between HXK1 and oxylipins has been described, it might be speculated that an HXK1-mediated re-allocation of carbon flux from cytosolic to plastidial glycolysis might also affect oxylipin metabolism through biosynthesis of PUFAs. Because the oxylipin OPDA (12-oxophytodienoic acid) is precursor of jasmonic acid (JA), this would add an additional layer of the complex interaction of HXK1 with phytohormones. ^*17*^, in their pioneering work on the role of HXK1 as a glucose sensor, demonstrated reduced sensitivity of the *gin2-1* mutants to auxins. Since both, JA and auxin, use shared components in their signaling cascade, JA is considered to reduce auxin responsiveness through recruitment of such shared components ^*18*^.

In addition to redox homeostasis, inverse dynamics of plastidial and mitochondrial pyruvate in *hxk1* may also impact fatty acid biosynthesis in chloroplasts. The proteome of this biosynthetic pathway revealed opposite trends for *hxk1* and wildtype of central enzymes during early eCO_2_. Plastidial pyruvate dehydrogenase (pPDH) E1 component (AT1G30120), 3-oxoacyl-[acyl-carrier-protein] synthase I (AT5G46290) and ATP-citrate synthase B-1 (AT3G06650) were upregulated in Col-0 but downregulated in *hxk1* under eCO_2_ (Figure 5 A-C). Also, carboxylic acid metabolism, for which pyruvate represents a central substrate, was affected in *hxk1* at the proteome level (Supplementary Figure S9). All together, these observations demonstrate that HXK1 reaches out far beyond cytosolic carbohydrate metabolism, influencing subcellular redox balance, fatty acid and lipid metabolism and may also integrate ROS metabolism with phytohormonal response.

### Metabolite dynamics in a heterozygous *bou* mutant point to a new function of this mitochondrial transporter

The design of the present study was not compatible with the use of the homozygous *bou* mutant, which does not grow at ambient CO_2_ levels used as starting conditions. Thus, we made use of heterozygous mutant plants, *h-bou*, which still have an 85% growth reduction as compared to the wild type (Supplementary Figure S10), but are able to develop mature leaves at ambient CO_2_ levels. As expected, glycine significantly accumulated in mitochondria of *h-bou* at aCO_2_, although not as pronounced as reported for homozygous mutants (Figure 6A). In line with glycine accumulation, the H subunit of the glycine cleavage system (AT2G35370) exhibited lower expression levels in *h-bou* and remained consistently lower than wildtype under all conditions (Figure 6B). Opposite trends were observed in expression of other subunits of the GDC system (AT4G33010 and AT2G26080; Figure 6 C, D) revealing a significant decrease in *h-bou*, but increased expression and a final stabilization in wildtype. Serine accumulated slowly in plastids of *h-bou* plants after transfer to eCO_2_ (Figure 7 A, B), pointing to a shift of serine production from photorespiratory to the plastidial phosphorylated pathway (PPSB). This was supported by a higher expression rate of SHMT3 (AT4G32520) in *h-bou* samples as compared to Col-0 (Figure 7 C), and also by glycine accumulation in plastids of *h-bou*.

**Figure 6.**
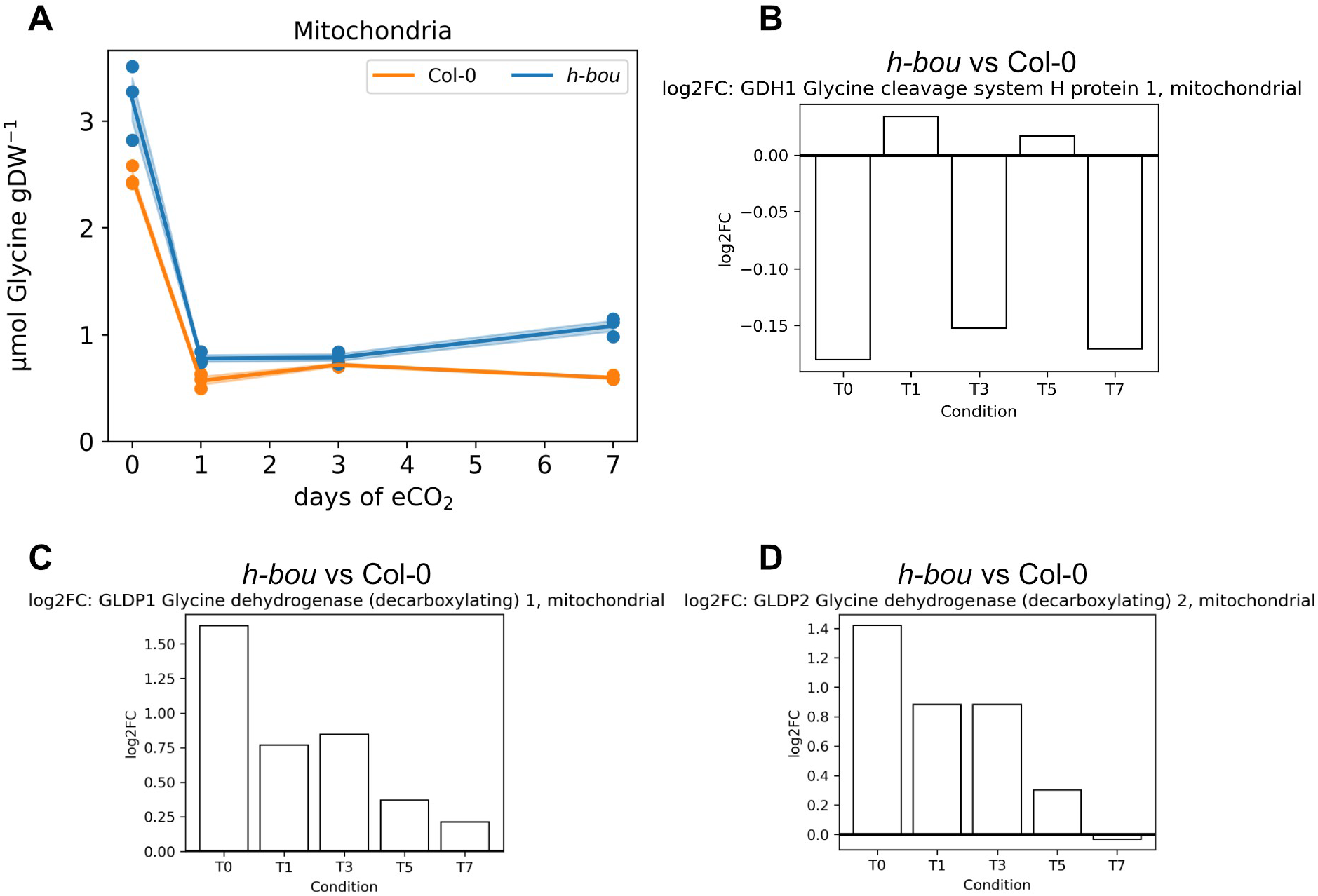
Mitochondrial glycine dynamics with transcripts of associated enzymes. (A) Dynamics of mitochondrial glycine concentrations in *h-bou* (blue) and Col-0 (orange). Ordinate reflects glycine amounts in µmol gDW^-1^, abscissa shows time of exposure to eCO2 in days. (B) Log2FC of GDC-H (AT2G35370) comparing *h-bou* to Col-0, (C) Log2FC of GDC-P1 (AT4G33010) comparing *h-bou* to Col-0– (D) Log2FC of GDC-P2 (AT2G26080) comparing *h-bou* to Col-0; n = 3.

**Figure 7.**
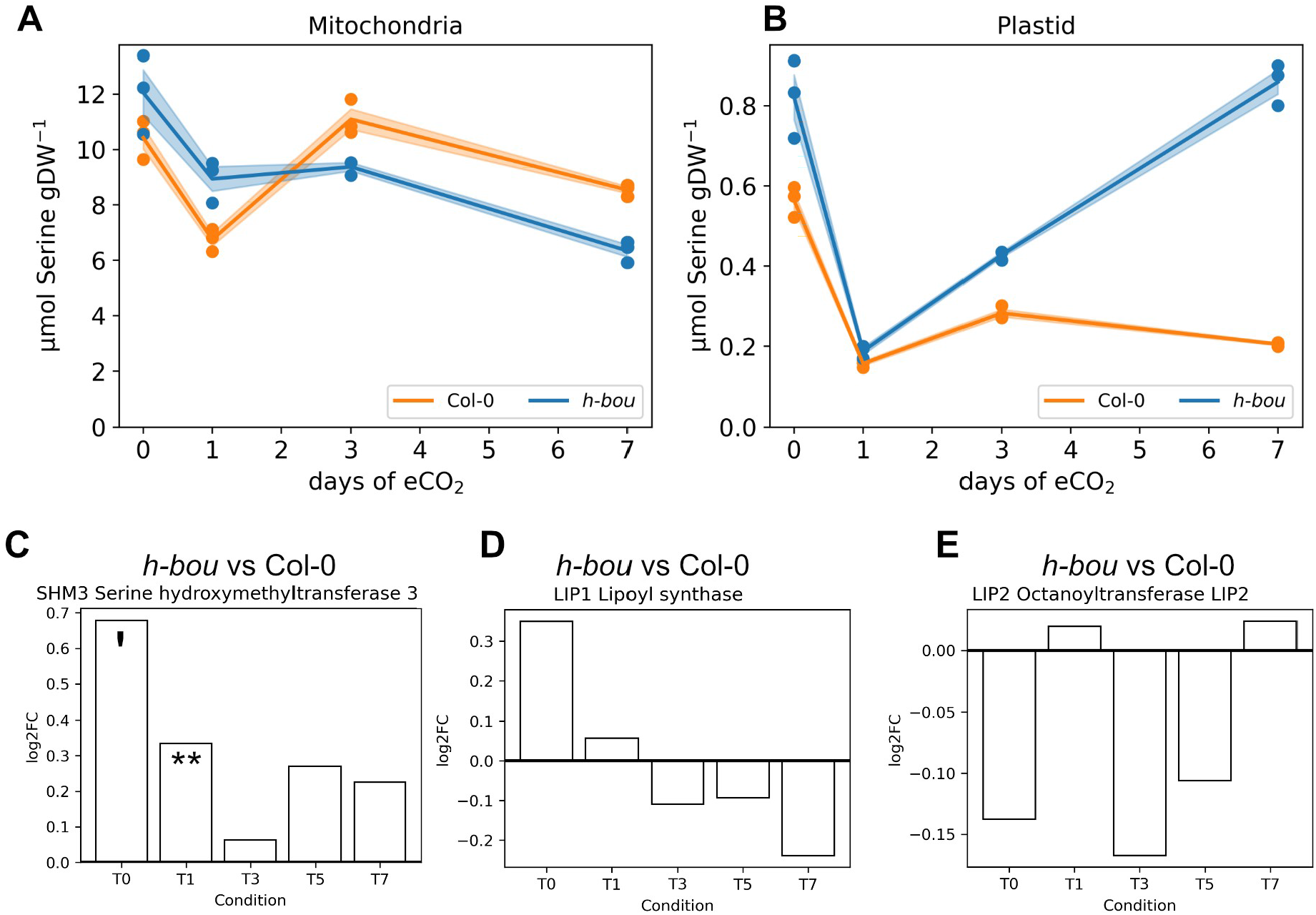
Subcellular serine dynamics in mitochondria and plastids, with transcripts of relevant enzymes. (A) Dynamics of mitochiondrial serine concentration in *h-bou* (blue) and Col-0 (orange), (B) Dynamics of plastidial serine concentration in *h-bou* (blue) and Col-0 (orange). Ordinates reflect serine amounts in µmol gDW^-1^, abscissae show time of exposure to eCO2 in days. (C) – (E) Log2FC between *h-bou* and Col-0 for SHMT3, LIP1 and LIP2, respectively (** - p value < 0.01, ‘ - p value < 0.1; n = 3 replicates).

Considering the reported function of BOU as a mitochondrial glutamate transporter necessary for polyglutamylation of THF ^*10*^, an elevated, not reduced, glutamate concentration in *h-bou* mitochondria at aCO_2_ was unexpected (Supplementary Figure 11 A). However, already in 2002, it was reported that the BOU protein belongs to the family of carnitine /acylcarnitine transporters ^*19*^ that transport fatty acid carnitine esters into the mitochondrial matrix, where they are involved in lipoylation of decarboxylase enzymes, including the glycine cleavage system ^*20*^. Although lipoic acid was not measured in this study, strong evidence for disturbed homeostasis in *h-bou* comes from significant differences in expression levels of transcripts involved in lipoic acid *de novo* synthesis, LIP1 (AT2G20860) and LIP2 (AT1G04640). Under ambient conditions, LIP1 showed higher expression in *h-bou*, and expression significantly dropped and fell below wild type levels upon transfer to eCO_2_ (Figure 7 D). There was no marked difference in the expression pattern of LIP2 between *h-bou* and the wild type, but the level of gene expression in *h-bou* was generally decreased (Figure 7 E).

If lipoate availability was reduced in mitochondria of *h-bou*, other mitochondrial decarboxylases featuring E2 subunits, e.g., PDH, 2OGDH and the branched-chain α-ketoacid dehydrogenase complex (BCKDC) involved in catabolism of branched chain amino acids, should be affected ^*21*^. Indeed, we observed elevated pyruvate levels in mitochondria of *h-bou* at aCO_2_, indicating reduced turnover (Figure 2B) despite a higher expression level of three different PDH subunits (AT1G48030, AT3G52200, AT1G59900), including the lipoate depending E2 subunit AT1G48030 (Supplementary Figure 12). Upon transition to eCO_2_, a significant decrease in expression of these subunits was observed, which, in conjunction with decreasing pyruvate levels, points to a more efficient turnover, possibly because of a released competition for lipoic acid between the glycine cleavage system and PDH when photorespiratory activity declined. A significant decrease in the transcript level of the E2 subunit of PDH (LTA3, AT3G52200) at eCO_2_, indicating return to normal TCA function, supports this view.

Expression of the E2 subunit of 2-oxoglutarate dehydrogenase, another consumer of lipoate, displayed an increasing trend after transition to eCO_2_ (Supplementary Figure S11 C, D), accompanied by an increase in its substrate 2-oxoglutarate, reaching wildtype levels at day 3 (Supplementary Figure S11 B). The low mitochondrial 2-oxoglutarate concentration at aCO_2_ indicated a disruption of normal TCA cycle function. Low conversion of 2-oxoglutarate to succinyl-CoA affords replenishment through anaplerotic reactions, an example being catabolism of the branched chain amino acids (BCAA) leucine, valine and isoleucine, all of which were significantly depleted in *h-bou* mitochondria under aCO_2_ and gradually accumulated upon transition to eCO_2_ (Figure 8, A-C). It is worth mentioning that BCAA catabolic reactions are also coupled to the transamination of 2-oxoglutarate to glutamate ^*22*^. The enzyme BCKDC, involved in catabolism of BCAA, is another consumer of lipoate, thus creating a complex network of competition, were elevated transcript levels of BCKDC (Figure 8 D) at aCO_2_ and its decline at eCO_2_ contribute to the view of disturbed lipoic acid homeostasis in the *h-bou* mutant. Interestingly, all three branched chain amino acids accumulated in the homozygous *bou* mutant when shifted from high to aCO_2_ ^*23*^. Under this condition, a high demand for GDC activity is created, thus restricting lipoate availability for other enzymes. ^*23*^ pointed out that the alteration in BCAA levels were not the result of low nitrate supply causing re-channeling of amino acids, and they also demonstrated that the E2 subunit of BCKDC was elevated at aCO_2_. While this is in line with the phenotype of homozygous *bou* being stronger than in the heterozygous condition, the question still remains, why mitochondrial Glu levels were even higher in *h-bou* than in the wildtype. One obvious explanation is the transamination of 2-oxoglutarate to glutamate in the course of BCAA catabolism needed to replenish succinyl-CoA ^*24, 25*^, which would also explain low 2-oxoglutarate. Another, though less likely, explanation would be a metabolic transition to the import and use of glutamate for 2-oxoglutarate production in the mitochondria in order to fill up the TCA cycle at limiting PDH activity. Both are compatible with limiting lipoate availability in mitochondria as a result of BOU dysfunction, and thus we conclude that the combined omics approach applied here gives strong evidence to a function of BOU as carnitine/acylcarnitne transporter.

**Figure 8.**
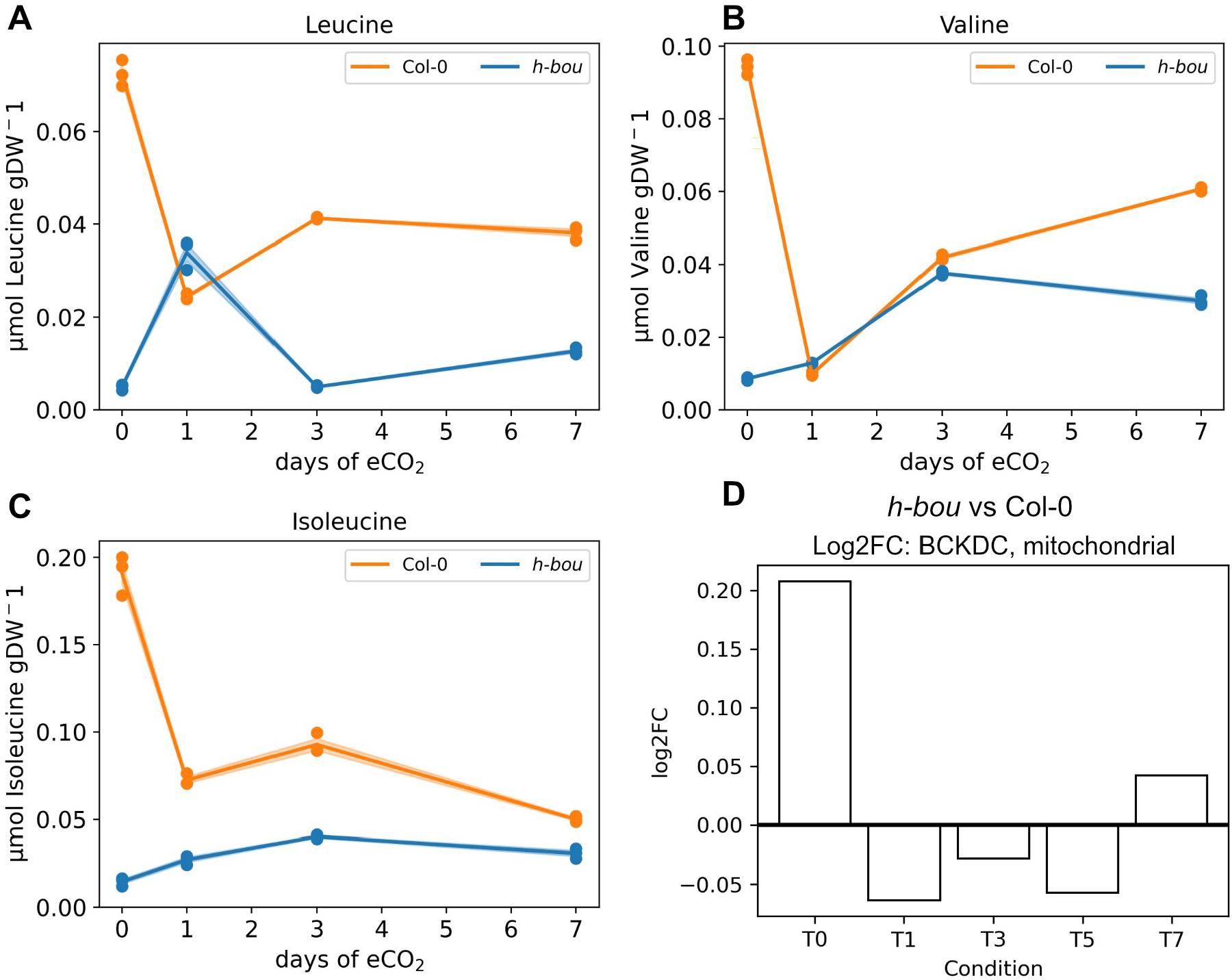
Mitochondrial concentrations of branched chain amino acids, with transcripts of catabolic enzyme BCKDC. (A) Dynamics of mitochiondrial leucine concentration in *h-bou* (blue) and Col-0 (orange), (B) Dynamics of mitochondrial valine concentration in *h-bou* (blue) and Col-0 (orange). (C) Dynamics of mitochondrial isoleucine concentration in *h-bou* (blue) and Col-0 (orange). Ordinates reflect amounts in µmol gDW^-1^, abscissae show time of exposure to eCO2 in days. (D) Log2FC between *h-bou* and Col-0 for BCKDC (AT3G06850); n = 3.

Taken together, our combined and subcellular resolved omics approach revealed additional, new information on complex effects of mutations and helps to elucidate non-obvious phenotypes. It clearly shows that metabolome analysis at the whole cell level reaches limits when it comes to reaction pathways like, e.g., photorespiration, that are shared among various cellular compartments.

## Material and Methods

### Plant material and growth condition

*Arabidopsis thaliana* (L.) Heynh. Col-0 as wild type, and mutants *hpr1-1* (SALK067724), *hxk1* (*gin2-1*, Salk_034233c, At4g29130) and heterozygous *bou* (GK-079D12.01, At5g46800) were used in this study. All plants were grown in fully controlled growth chambers at Stuttgart University for six weeks in soil (seedling substrate, Klasmann-Deilmann GmbH) under ambient carbon dioxide (450 ± 20 ppm) and short day (8 h/16 light/dark, 100 µmol m^-2^ s^-1^, 60% relative humidity, temperature 22/16 °C). Standard NPK fertilizer was applied immediately after thinning seedlings and then every 2 weeks until harvest. After 6 weeks, one set of plants (9 per genotype) was sampled (day 0), and the rest transferred to elevated CO_2_ (1000 ± 20 ppm) at the same environmental settings. Consecutive sampling was done at days 1, 3, 5 and 7, each time 4 h into the light phase at the middle of a short day. For each genotype, three biological replicates were harvested, each consisting of a pool of three full rosettes (3 replicates per each genotype and time points, totally 60 samples). Aliquots of fresh plant material were prepared for Proteomics and Transcriptomics analysis, and the rest subjected to lyophilization (Heto PowerDry LL3000; Thermo Electron C.) for subcellular fractionation.

### RNA-seq analysis

Between 16 and 24 million, 100 bp, paired-end reads per sample were quality checked using FastQC (https://www.bioinformatics.babraham.ac.uk/projects/fastqc/), and adapter trimming or low-quality filtering were done using Trimmomatic ^*26*^. Filtered reads were mapped to the Arabidopsis genome assembly with Tophat2 ^*27*^. Counting and normalization of mapped reads and analysis of differential expression were done using the cufflinks suite ^*28*^.

### Proteome analysis by mass spectrometry

Protein extraction and trypsin digestion were carried out following the protocol of (Marino et al. 2019). Liquid chromatography-tandem mass spectrometry (LC-MS/MS) was conducted as previously described, with peptides separated over a 90-minute linear gradient of 5–80% (v/v) CAN ^*29*^. Raw data files were processed using MaxQuant software version 2.4.14.0 ^*30*^. Peak lists were searched against the Arabidopsis reference proteome (Uniprot, www.uniprot.org) using default settings, with the ‘match-between-runs’ feature enabled. Protein quantification was performed using the label-free quantification (LFQ) algorithm ^*30*^. Subsequent analysis was executed using Perseus version 2.0.11 ^*31*^. Potential contaminants, proteins identified only by site modification, and reverse hits were excluded from further analysis. Only protein groups quantified by the LFQ algorithm in at least three out of four replicates in at least one condition were considered. LFQ intensities were log2-transformed, and missing values were imputed from a normal distribution using Perseus with standard settings.

### Nonaqueous fractionation

Subcellular fractionation of lyophilized plant samples by NAF procedures was performed as earlier described ^*32, 33*^. Briefly, approximately 80-100 mg of lyophilized leaf homogenate were suspended in 10 ml of ice-cold mixture of tetrachloroethylene-heptane (solvent A, ρ = 1.36 g cm^-3^) and sonified on ice bath for intervals of 5 s with 15 s pauses over a total time course of 12 min (Branson Sonifier 250, Branson, USA; output control 3). Under constant cooling, sonified homogenate was sieved through nylon gauze, pore size 30 µm, and the filtrate was centrifuged for 10 min at 2350 g, 4 °C Supernatant was discarded and pellet resuspended in 1.5 ml of fresh cold solvent A, subsequently loaded onto the ice-cold non-aqueous linear gradient combination of organic solvents initiated by solvent A (ρ = 1.36 g cm^-3^), and ending up to pure tetrachloroethylene (ρ = 1.6 g cm^-3^). Gradients were subjected to ultracentrifugation (Optima™ L-90K, BeckMan Coulter, Ireland) for 3 h at 121,000 g, 4 °C. Fractionation of centrifuged gradient was performed into nine 1 ml fractions and each fraction aliquoted to 5 equal subfractions and immediately dried under vacuum condition for subsequent metabolite and marker enzyme analysis. Alkaline pyrophosphatase served as plastidial marker, UGPase as cytosolic marker, succinyl-CoA-synthetase as mitochondrial marker, and acid phosphatase as vacuolar marker enzyme as described earlier ^*32-34*^.

### Metabolic profiling

For metabolite analysis, one aliquot of NAF and a respective whole cell sample was subjected to carbohydrate determination by HPLC (Dionex ICS 6000, ThermoFisher Scientific, USA), yielding concentrations of glucose, fructose, sucrose and raffinose as described by ^*35*^. Briefly, extraction was performed in 80% ethanol at 80 °C, followed by vacuum drying. Dried extracts were resuspended in Honeywell chromatographic grade water and subjected to HPLC analysis on Dionex CarboPac PA1 BioLC column (4 × 250 mm, ThermoFisher Scientific, USA). Carboxylic acids, including pyruvate, malate, fumarate and citrate as well as minerals (nitrate, phosphate, sulfate) were quantified by anion-exchange chromatography using another aliquot of samples as described ^*35*^. Briefly, extraction was done in 1 ml of 55 °C Honeywell water and incubation at 95 °C, followed by separation on Dionex IonPac AS11-HC RFIC column (4 × 250 mm, ThermoFisher Scientific, USA).

Amino acid measurements were performed by quantitative GC-MS/MS as described ^*36-38*^ with a few modifications using a third sample aliquot. Basically, a modified solid phase extraction of amino acids was performed by suspending NAF samples in 1 ml of 10 mM HCl and 10 min shaking at RT, followed by 2 min centrifugation at 14,000 g, RT. Hundred microliter of supernatant in addition to 10 nmol of norvaline (Acros Organics) as internal standard were subjected to amino acid purification, using homogenized suspension of 100 mg ml^-1^ of ion-exchange resin (DOWEX 50WX4, 200-400 mesh) in 10 mM HCl, incubated for 15 min at RT in Mobicol spin classic tubes equipped with 10 µm pore sized filters (MoBiTec GmbH, Germany), followed by centrifugation 5 min at 1,000 g, RT. Resin was washed twice by 80% methanol and 1 min centrifugation each time at 1,000 g, RT to remove non-amino acid metabolites. Subsequently, resin containing purified amino acids was suspended in 150 µl of 1:1 mixture of methanol and 8 M ammonia and centrifuged for 2 min at 5,000 g, RT. Eluent containing purified amino acids was dried in a speed vacuum concentrator (ScanSpeed 32, Denmark). Upon drying, derivatization was done by adding 50 µl of both MTBSTFA (Sigma Aldrich) and acetonitrile, 1 h incubation at 95 °C followed by 2 h at RT and subsequent analysis by GC-MS/MS (TQ8040, Shimadzu, Japan). One microliter of the derivatized samples was injected to device, applying helium as carrier gas at a flow of 1.12 ml/min. Stationary phase was a 30 m Optima 5MS-0.25 μm fused silica capillary column. Ion source, column oven and injection temperature were set up at 250 °C, 100°C and 250 °C respectively. A split 10 gradient program was applied with initial column temperature of 100 °C for 1 min, followed by 15 °C increment per minute till 290 °C, holding for 3 min, again followed by same increment rate to reach the final temperature as 330 °C and holding for 10 min. Subsequent to 5 min solvent delay, spectra of MS device were recorded in Q3 scanning mode.

All other metabolites including carbohydrates, sugar alcohols, carboxylic acids and polyamines were measured using the last remaining sample aliquots by GC-MS/MS as described earlier ^*39*^. Briefly, extraction on ice into methanol-chloroform-water mixture, 2.5:1:0.5 (v/v/v) was followed by centrifugation. Polar phase was separated and along with norvaline as internal standard, dried in a speed vacuum concentrator equipped with cold trap (ScanVac, Denmark). After drying, methoximation was performed using 20 µl of methoxamine dissolved in pyridine (40 mg ml^-1^) by 90 min incubation at 30 °C followed by silylation, using 80 µl of MSTFA (Sigma Aldrich) and 30 min incubation at 50 °C. Detection was done by the same GC-MS/MS device as amino acids and a different split 10 gradient program as 70 °C for initial temperature and one min holding, followed by 15 °C increment per minute till reaching to final 330 °C, and holding for 10 min. Ion source, column oven and injection temperature were set up at 250 °C, 70°C and 230 °C, respectively, and after 4.7 min solvent delay, spectra were recorded in Q3 scanning mode.

### OmicsDB Tech Stack

In order to quickly access omics data in a reliable and reproducible manner, an SQLite Database (OmicsDB) was created, which can be accessed via a custom API (OmicsAPI) in python (Version 3.11.5). When utilizing this API in order to submit data to the OmicsDB, the API sends a URL request to the API of Panther knowledgebase (PANTHER 18.0) in order to retrieve the associated GO terms of a transcript or a protein ^*40*^. In addition to this, the organism-specific locus name is gathered via the API of the UniProt knowledgebase (Release 2024_01) ^*41*^. In order to associate proteins and transcripts to metabolites, GO terms where matched to KEGG modules (Release 108.1), from which information on metabolites was obtained via the associated API ^*42-44*^. The data stored in the OmicsDB can now be accessed via the OmicsAPI in a Go Term specific manner, e.g. “Photorespiration” or “Chloroplast”. For fast and easy data exploration, an additional analysis API (ExplorerAPI) was created which can perform principle component analysis (PCA), partial least squares regression (PLS) between Datasets, and time-lagged cross correlation analysis (TLCC). This API should be used in a Jupyter Notebook. Both PCA and PLS can be performed in a sparse manner and use scikit-learn (Version 1.3.2) or in the case of sPLS, the R package mixOmics (Version 6.22.0) in combination with rpy2 (Version 3.5.15) in order to make it accessible in python. Methods related to the TLCC were coded inhouse. In Addition, the ExplorerAPI provides basic plotting methods for individual Omics samples.

The entire codebase and database are available under https://git.nfdi4plants.org/thomas.naegele/OMICS_DB. Additional requirements regarding packages and package versions can be also viewed on GitHub.

The metabolite list associated with photorespiration was further extended to include glutamine, aspartate, malate, and asparagine.

### Statistics

Statistics were performed in python (Version 3.11.5). In order to calculate significance of log2FC, z-tests were performed after calculating the geometric mean and the propagation of error as suggested in ^*45*^. In addition, the p-value was adjusted for multiple testing using Bonferroni correction. The script was coded in house, in order to integrate with the OmicsAPI and is available on the same GitHub page.

## Acknowledgement

Prof. Hermann Bauwe and Dr. Stefan Timm (University of Rostock, Germany) are acknowledged for a generous gift of *hpr1-1* mutant seeds. We thank Annika Allinger and Laura Merkle for expert plant cultivation.

This work was supported by the German Science Foundation (DFG), HE3087/12-01 to AGH and NA1545/4-1 to TN.

## Author contributions

TN, AGH and DW designed the study. DS and JSH performed the experiments and analyzed the data. JG, EZ and JA performed transcriptome analysis. SS and DL performed and supervised proteomics analysis. All authors contributed to data evaluation and manuscript preparation. All authors read the manuscript and consent to its content.

## Data availability statement

Original data is available at https://git.nfdi4plants.org/thomas.naegele/Subcellular_analysis_of_metabolic_network_dynamics_under_elevated_CO2

## Supplementary Figures

**Supplementary Figure S1**. Principle component analysis of metabolites associated with photorespiration after exposure to elevated CO_2_. (A) Scatter plot of scores. The color of points indicates the genotype (Blue – *h-bou*, Orange – Col-0, Green – *hpr1-1*, Red -*hxk-1)*, while the shape indicates days of elevated CO_2_ (1000 ppm) treatment (Circle – T0, Triangle – T1, Square – T3, Pentagon – T7). T0 represents ambient conditions (approx. 400 ppm) before transfer to eCO_2_. Replicates: n = 3. (B) Metabolites showing the highest loadings, i.e., contribution to the principle components. The cut off for this table was set to 80% of the highest loading of each component. Loadings are sorted in descending order.

**Supplementary Figure S2**. Subcellular glycolate concentration and associated transporters. (A) Dynamics of glycolic acid in plastids, (B) Dynamics of glycolic acid in the cytosol. Ordinates reflect glycolic acid amounts in µmol gDW ^-1^ (means ± SE; n = 3), abscissae show time of exposure to eCO_2_ in days. (C) Log2FC of PLGG1 between *hpr1-1* and Col-0, (D) log2FC of BASS6 between *hpr1-1* and Col-0 (* - p value < 0.05, n = 3).

**Supplementary Figure S3**. Hypothetical bypass of HPR1. Expression data is given in log2FC between *hpr1-1* and Col-0. Order of squares represent order of conditions (1: aCO_2_, 2: 1 day eCO_2_, 3: 3 days eCO_2_, 4: 5 days eCO_2_, 5: 7 days eCO_2_). The top row represents transcriptomics data, while the bottom row represents proteomics data (* - p-val < 0.05; n = 3).

**Supplementary Figure S4**. Model of a data-derived suggested cytosolic bypass in *hpr1-1*.

**Supplementary Figure S5**. log2FC of PDK between *hpr-1* and Col-0 (*** - p value < 0.0001; n = 3).

**Supplementary Figure S6**. Cytosolic citrate dynamics (A) and Log2FC between *hpr1-1* and Col-0 of enzymes involved in citrate metabolism (means ± SE; n = 3). (B) Log2FC between *hpr1-1* and Col-0 of citrate synthase subunits. (C) Log2FC between *hpr1-1* and Col-0 of acetyl coenzyme A synthase and acetate/butyrate CoA ligase.

**Supplementary Figure S7**. Principal component analysis of eCO_2_ response in the proteomes of TCA cycle and cellular redox homeostasis. **(A)** Col-0, **(B)** *hxk1*, **(C)** *hpr1-1*, **(D)** *h-bou*. Colors and shapes indicate different time points of eCO_2_ exposure. Red filled circles: 0 days at eCO_2_, i.e., aCO_2_; olive filled triangles: 1 day at eCO_2_; green filled squares: 3 days at eCO_2_; blue cross: 5 days at eCO_2_; magenta crossed square: 7 days at eCO_2_. Data was scaled (z-scale, i.e., zero mean, unit variance).

**Supplementary Figure S8**. Dynamics of peroxisomal CATALASE2 (CAT2, AT4G35090) under eCO_2_. **(A)** Col-0, (**B**) *hxk1*, (**C**) *hpr1-1*, (**D**) *h-bou*.

**Supplementary Figure S9**. Principal component analysis of proteins involved in carboxylic acid metabolism. (A) PCA without loadings; (B) PCA with loadings. Colours indicate genotypes (red: Col-0; cyan: *hxk1*). Symbols represent duration of eCO_2_ treatment.

**Supplementary Figure S10**. Phenotype of the homozygous *bou* mutant **(A)**, the heterozygous mutant *h-bou* **(B)** and the Col-0 wildtype (**C)**. Plants were grown for six weeks in soil under ambient carbon dioxide (450 ± 20 ppm) and short day (8 h/16 light/dark, 100 µmol m-2 s-1, 60% relative humidity, temperature 22/16 °C) before photography.

**Supplementary Figure S11**. Mitochondrial glutamate and oxoglutarate dynamics with transcripts of associated enzymes. (A) Dynamics of mitochondrial glutamate concentrations in *h-bou* (blue) and Col-0 (orange). Ordinate reflects glutamate amounts in μmol gDW^-1^ (means ± SE; n = 3), abscissa shows time of exposure to eCO_2_ in days. (B) Dynamics of mitochondrial oxoglutarate concentrations in *h-bou* (blue) and Col-0 (orange). Ordinate reflects 2-oxoglutarate amounts in μmol gDW^-1^ (means ± SE; n = 3), abscissa shows time of exposure to eCO_2_ in days. (C) Log2FC of OGDH-E2 (AT4G26910) comparing *h-bou* to Col-0– (D) Log2FC of OGDH2-E2 (AT5G55070) comparing *h-bou* to Col-0; n = 3.

**Supplementary Figure S12**. Log2FCs of PDH subunits for *h-bou* as compared to Col-0 enzymes. (A) Log2FC between *h-bou* and Col-0 for the lipoate containing E2 subunit (AT1G48030). (B) Log2FC between *h-bou* and Col-0 for subunit 2-1 (AT3G52200). (C) Log2FC between *h-bou* and Col-0 for the E1 subunit (AT1G59900); n = 3.

