## Supplemental material for "Regulation of plant metabolism under elevated CO_2_"

### Supplements

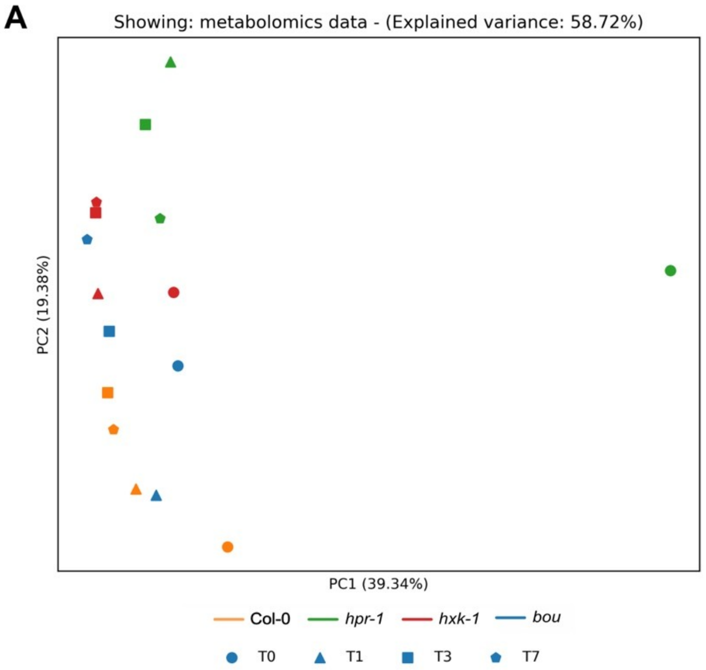


**B**

| Loadings PC1 | metabolite | compartment |
|---|---|---|
| 0.260 | Glycine | vacuole |
| 0.260 | Glycine | mitochondria |
| 0.260 | Glycine | cytosol |
| 0.253 | Glyceric acid | mitochondria |
| 0.251 | Serine | mitochondria |
| 0.251 | Serine | vacuole |
| 0.248 | Serine | cytosol |
| 0.246 | Serine | plastid |
| 0.230 | Glyceric acid | plastid |
| 0.230 | Malate | cytosol |
| 0.229 | Glyceric acid | vacuole |
| 0.219 | Glycine | plastid |

| Loadings PC2 | metabolite | compartment |
|---|---|---|
| 0.337 | Asparagine | plastid |
| 0.286 | Glutamine | mitochondria |
| 0.276 | Glutamine | cytosol |

**Supplementary Figure S1.** Principle component analysis of metabolites associated with photorespiration after exposure to elevated $CO_2$. (A) Scatter plot of scores. The color of points indicates the genotype (Blue – *h-bou*, Orange – Col-0, Green – *hpr1-1*, Red -*hxk-1)*, while the shape indicates days of elevated $CO_2$ (1000 ppm) treatment (Circle – T0, Triangle – T1, Square – T3, Pentagon – T7). T0 represents ambient conditions (approx. 400 ppm) before transfer to $eCO_2$. Replicates: n = 3. (B) Metabolites showing the highest loadings, i.e., contribution to the principle components. The cut off for this table was set to 80% of the highest loading of each component. Loadings are sorted in descending order.

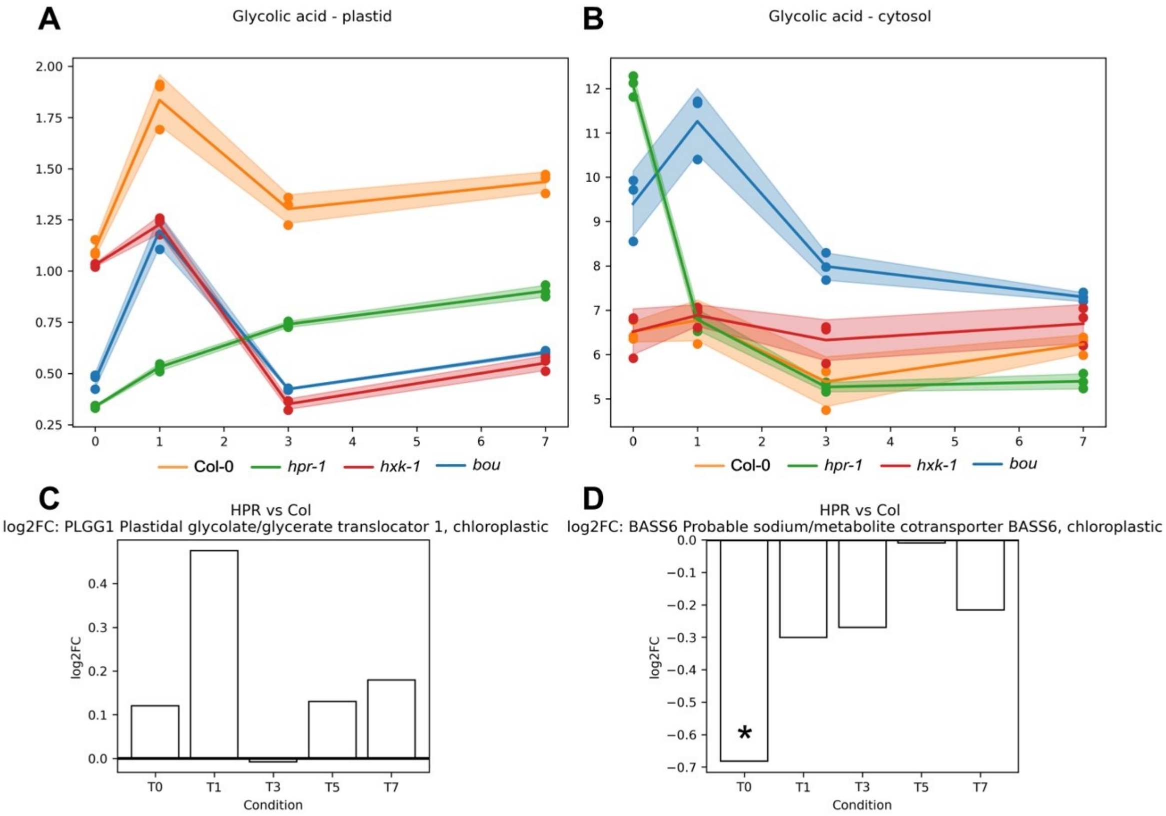


**Supplementary Figure S2**. Subcellular glycolate concentration and associated transporters. (A) Dynamics of glycolic acid in plastids, (B) Dynamics of glycolic acid in the cytosol. Ordinates reflect glycolic acid amounts in µmol $gDW^{-1}$, abscissae show time of exposure to $eCO_2$ in days. (C) Log2FC of PLGG1 between *hpr1-1* and Col-0, (D) log2FC of BASS6 between *hpr1-1* and Col-0 (* - p value < 0.05).

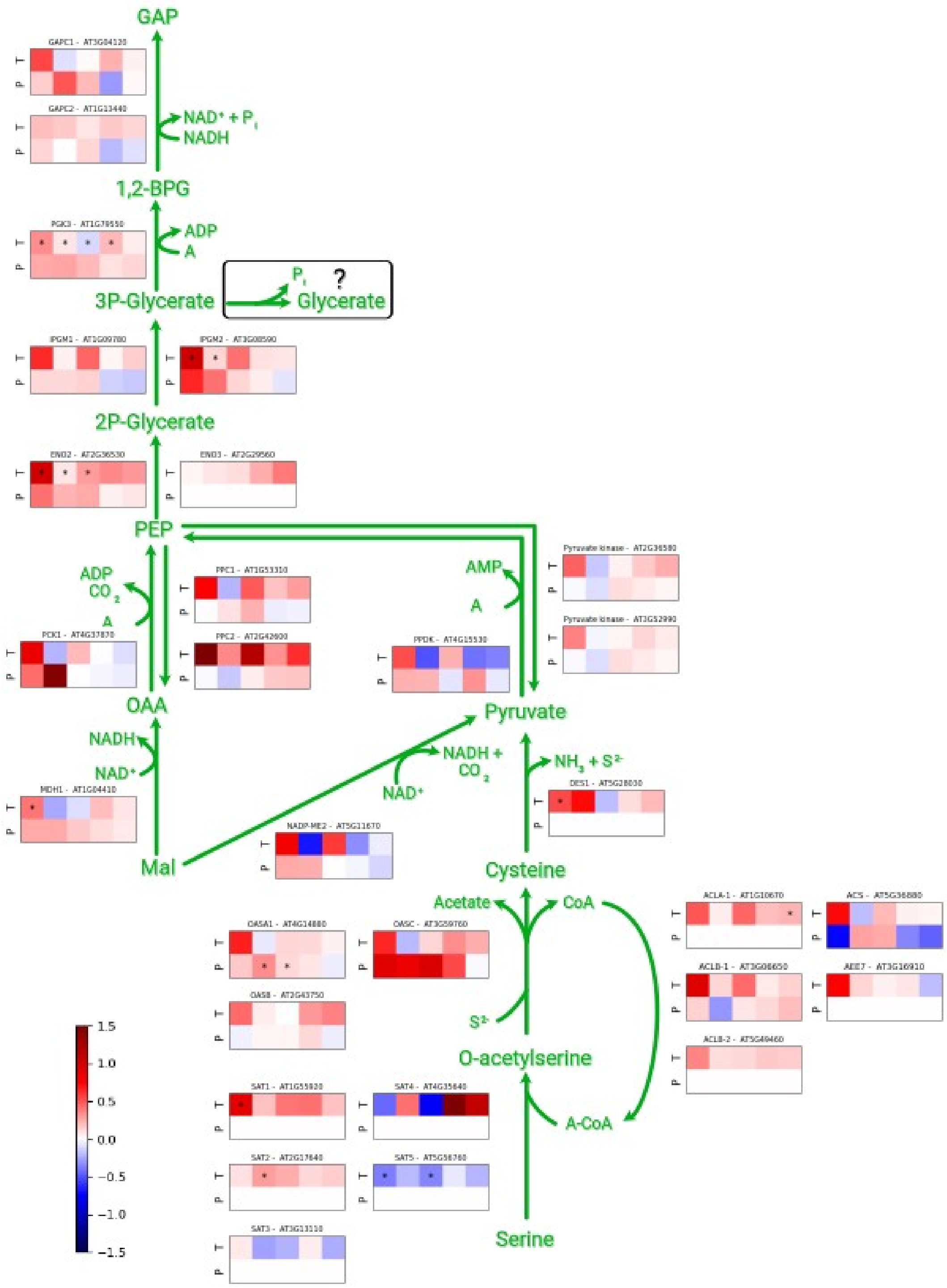


**Supplementary Figure S3.** Hypothetical bypass of HPR1. Expression data is given in log2FC between *hpr1-1* and Col-0. Order of squares represent order of conditions (1: a$CO_2$, 2: 1 day e$CO_2$, 3: 3 days e$CO_2$, 4: 5 days e$CO_2$, 5: 7 days e$CO_2$). The top row represents transcriptomics data, while the bottom row represents proteomics data (* - p-val < 0.05; n = 3).

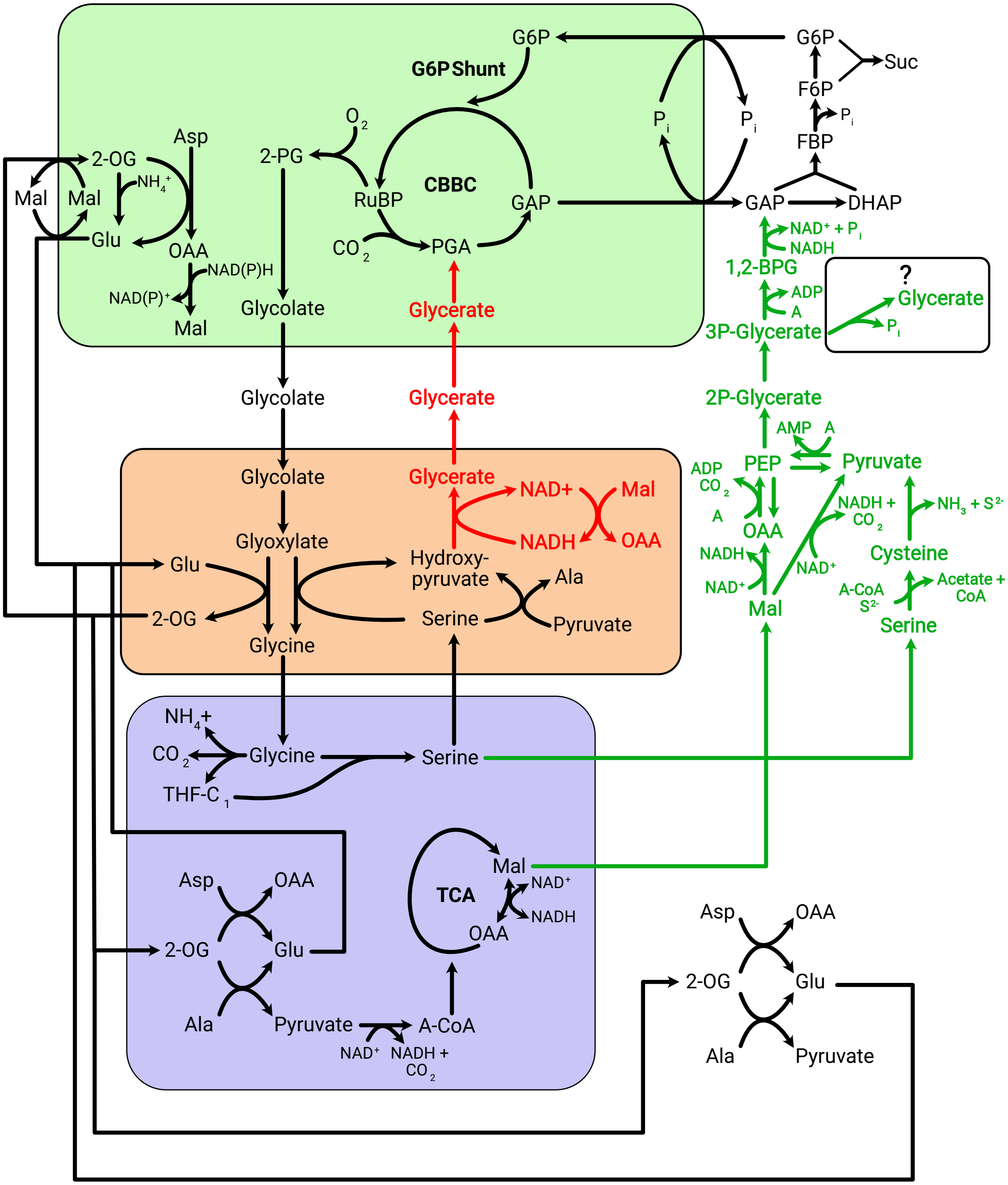


**Supplementary Figure S4.** Model of a data-derived suggested cytosolic bypass in *hpr1-1*.

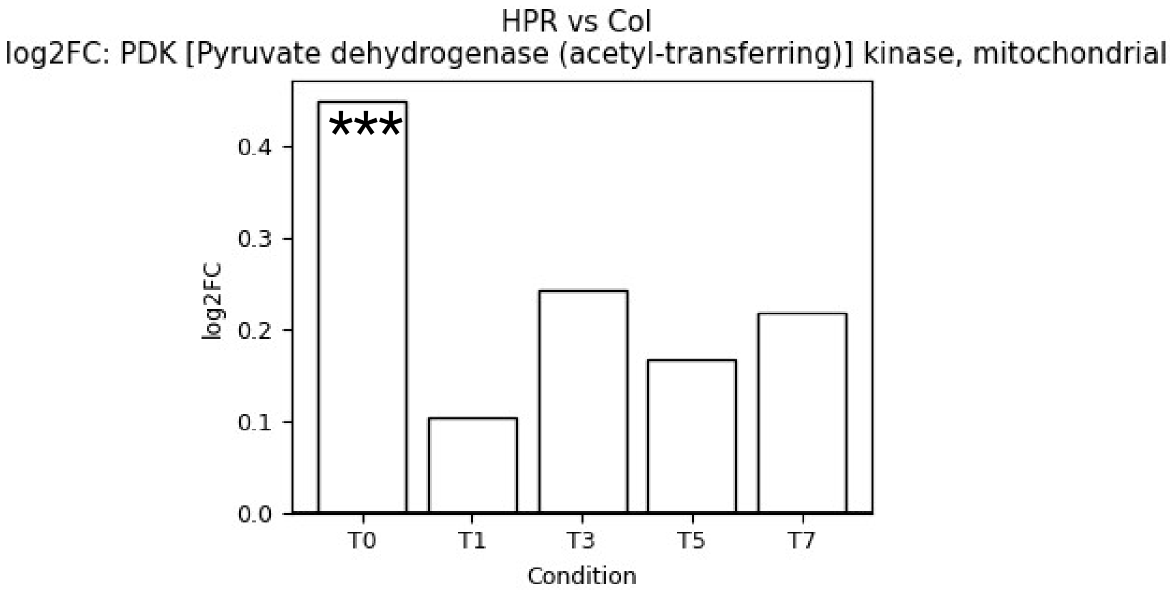


**Supplementary Figure S5**. log2FC of PDK between *hpr1-1* and Col-0 (*** - p value < 0.0001; n = 3).

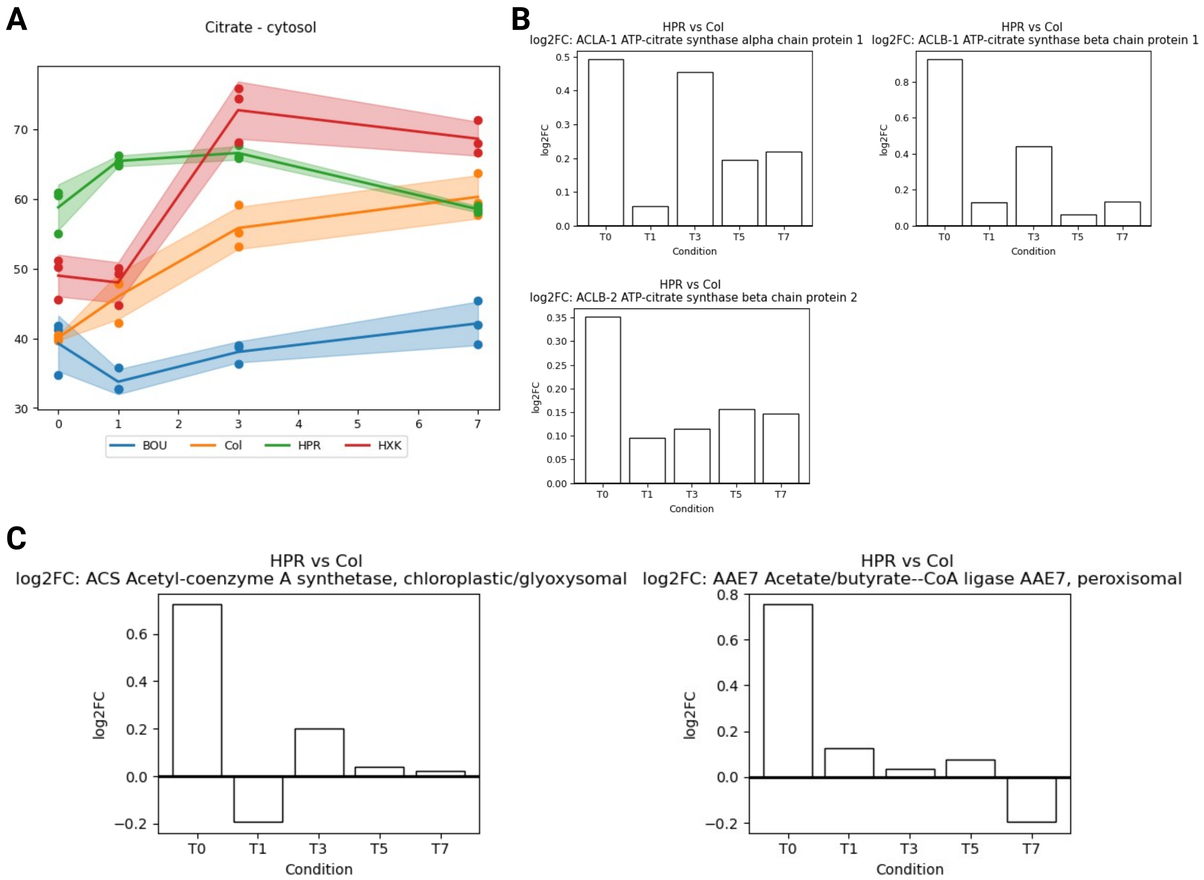


**Supplementary Figure S6.** Cytosolic citrate dynamics (A) and Log2FC between *hpr1-1* and Col-0 of enzymes involved in citrate metabolism (means ± SE; n = 3). (B) Log2FC between *hpr1-1* and Col-0 of citrate synthase subunits. (C) Log2FC between *hpr1-1* and Col-0 of acetyl coenzyme A synthase and acetate/butyrate CoA ligase.

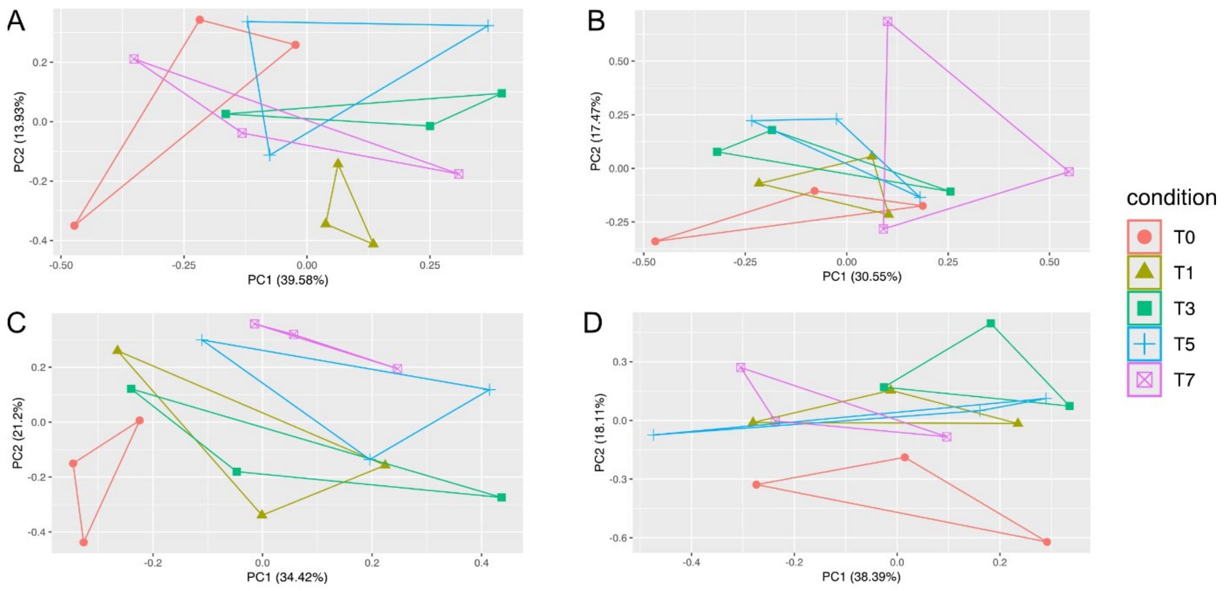


**Supplementary Figure S7.** Principal component analysis of e$CO_2$ response in the proteomes of TCA cycle and cellular redox homeostasis. **(A)** Col-0, **(B)** *hxk1*, **(C)** *hpr1-1*, **(D)** *h-bou*. Colors and shapes indicate different time points of e$CO_2$ exposure. Red filled circles: 0 days at e$CO_2$, i.e., a$CO_2$; olive filled triangles: 1 day at e$CO_2$; green filled squares: 3 days at e$CO_2$; blue cross: 5 days at e$CO_2$; magenta crossed square: 7 days at e$CO_2$. Data was scaled (z-scale, i.e., zero mean, unit variance).

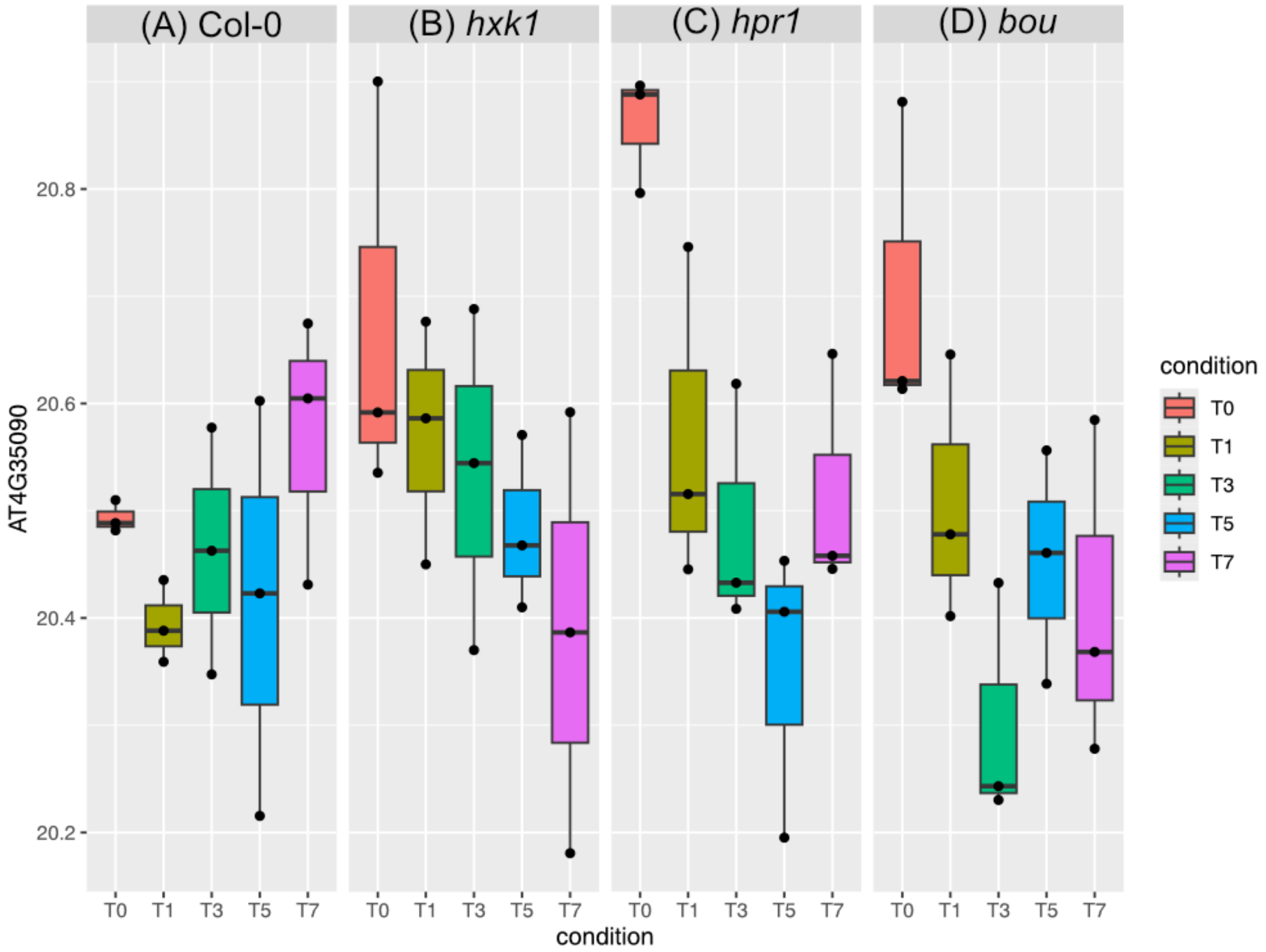


**Supplementary Figure S8**. Dynamics of peroxisomal CATALASE2 (CAT2, AT4G35090) under $eCO_2$. **(A)** Col-0**,** (**B**) *hxk1*, (**C**) *hpr1-1*, (**D**) *h-bou*.

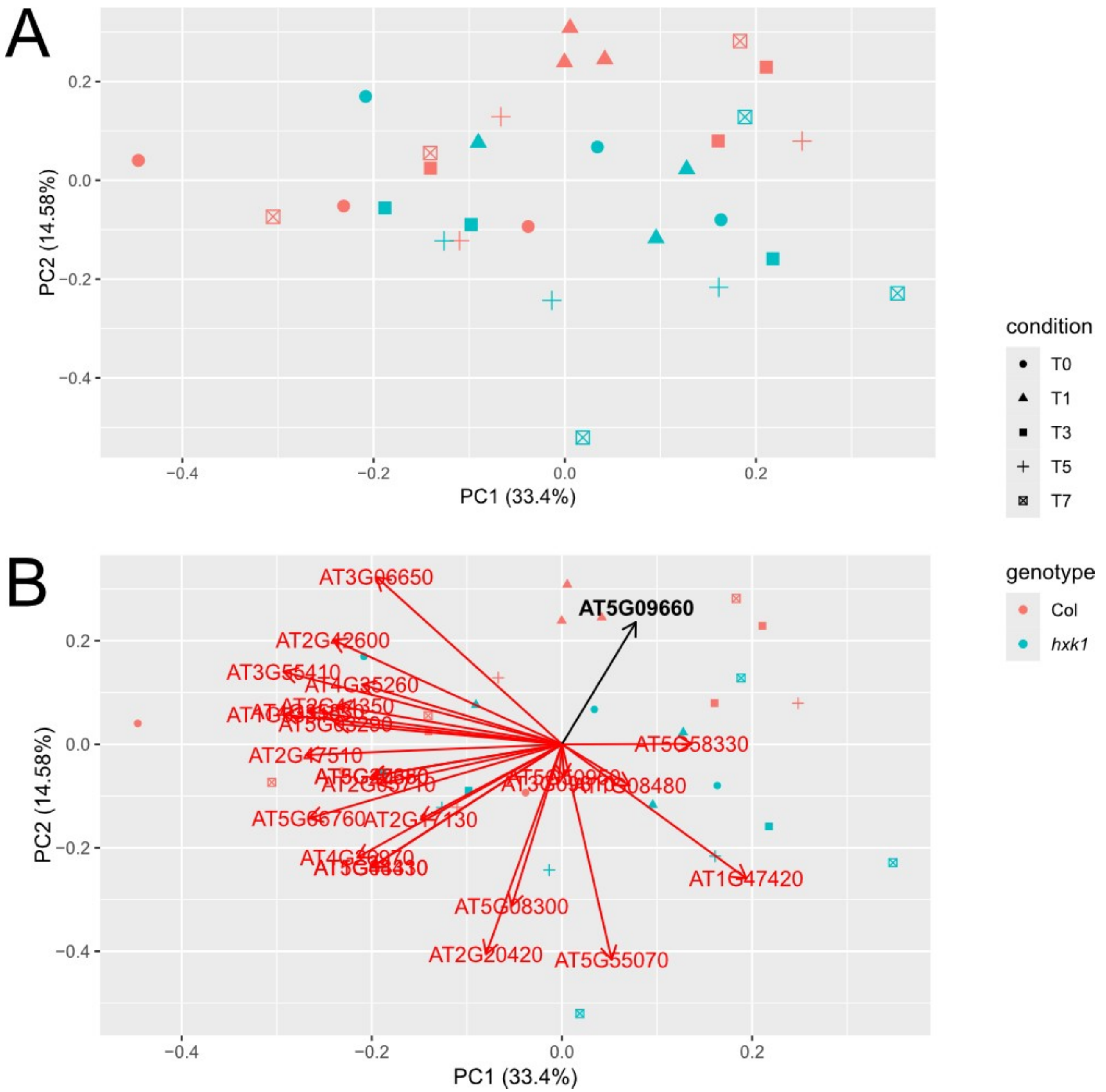


**Supplementary Figure S9.** Principal component analysis of proteins involved in carboxylic acid metabolism. (A) PCA without loadings; (B) PCA with loadings. Colours indicate genotypes (red: Col-0; cyan: *hxk1*). Symbols represent duration of $eCO_2$ treatment.

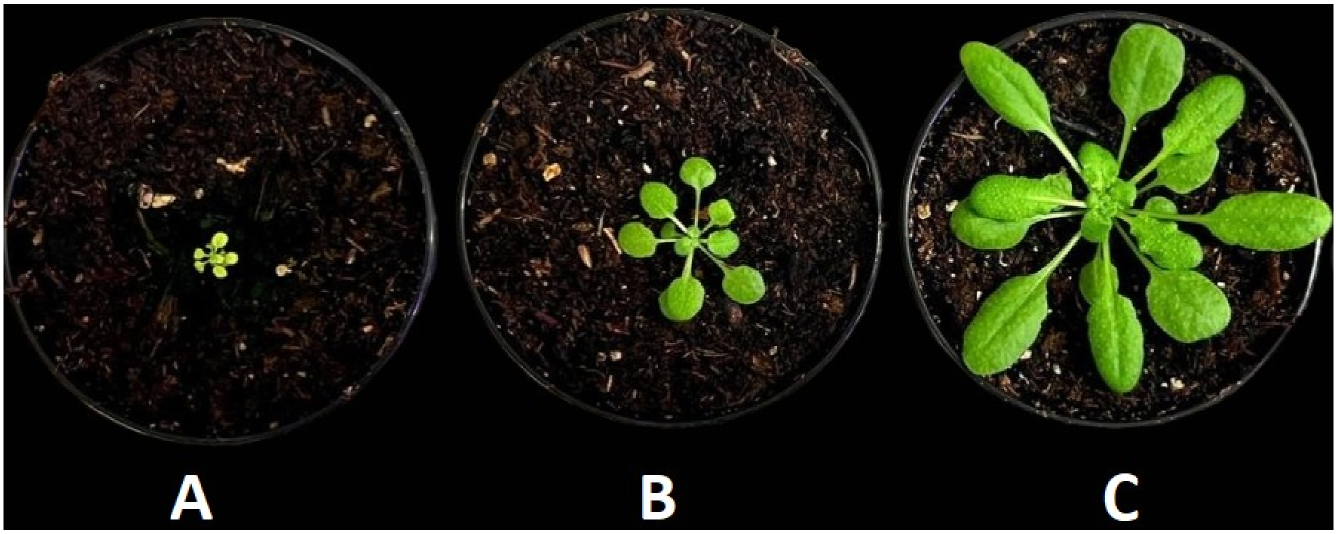


**Supplementary Figure S10.** Phenotype of the homozygous *bou* mutant **(A)**, the heterozygous mutant *h-bou* **(B)** and the Col-0 wildtype **(C).** Plants were grown for six weeks in soil under ambient carbon dioxide (450 ± 20 ppm) and short day (8 h/16 light/dark, 100 µmol m-2 s-1, 60% relative humidity, temperature 22/16 °C) before photography.

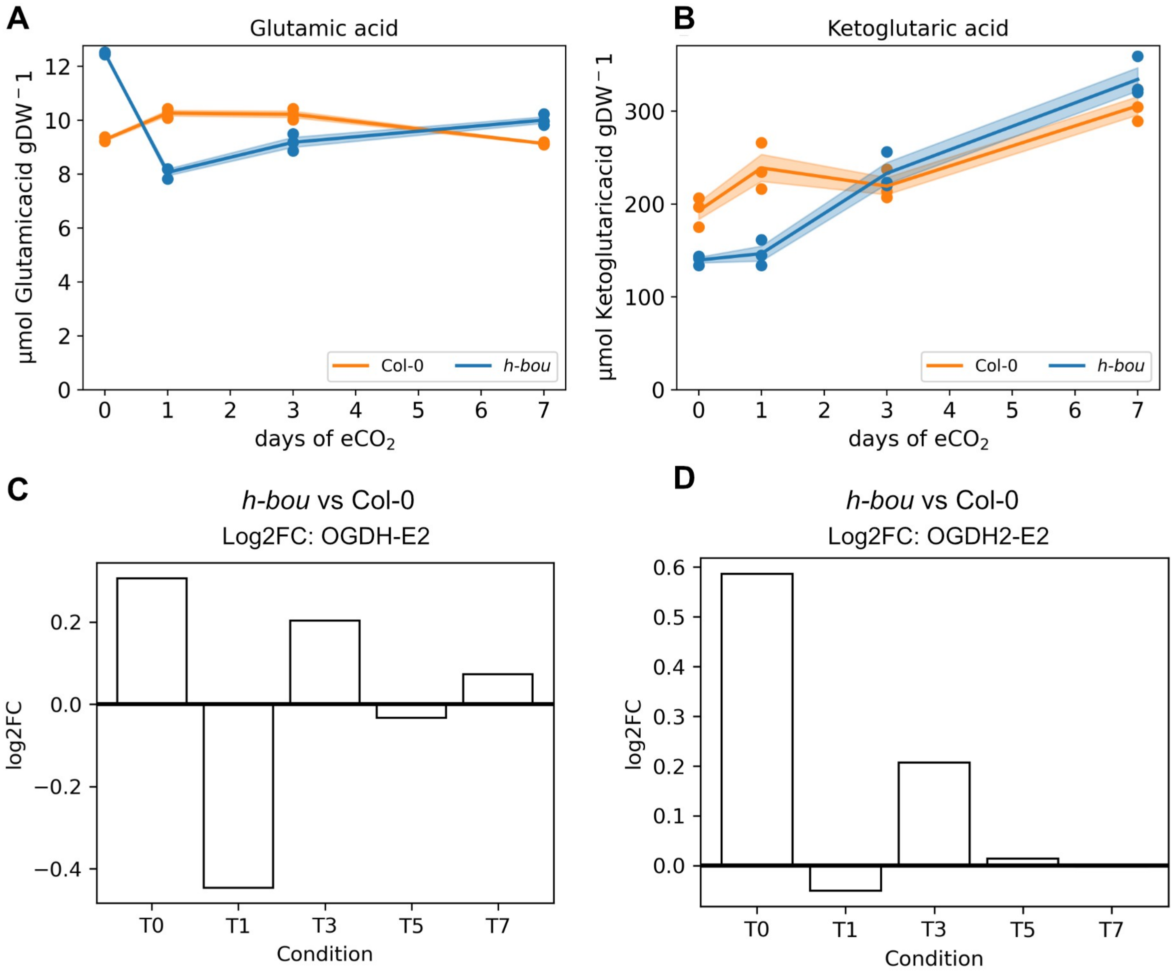


**Supplementary Figure S11.** Mitochondrial glutamate and oxoglutarate dynamics with transcripts of associated enzymes. (A) Dynamics of mitochondrial glutamate concentrations in *h-bou* (blue) and Col-0 (orange). Ordinate reflects glutamate amounts in µmol $gDW^{-1}$ (means ± SE; n = 3), abscissa shows time of exposure to $eCO_2$ in days. (B) Dynamics of mitochondrial oxoglutarate concentrations in *h-bou* (blue) and Col-0 (orange). Ordinate reflects 2-oxoglutarate amounts in µmol $gDW^{-1}$ (means ± SE; n = 3), abscissa shows time of exposure to $eCO_2$ in days. (C) Log2FC of OGDH-E2 (AT4G26910) comparing *h-bou* to Col-0– (D) Log2FC of OGDH2-E2 (AT5G55070) comparing *h-bou* to Col-0; n = 3.

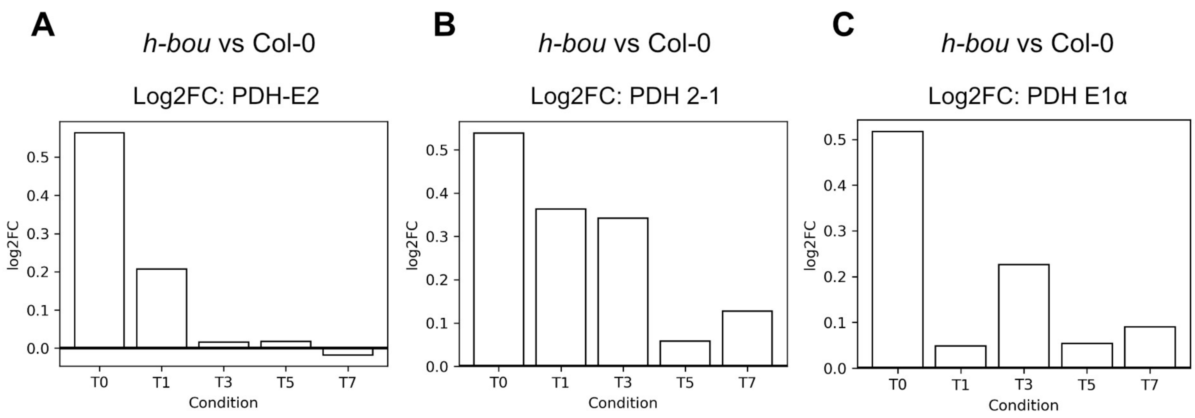


**Supplementary Figure S12.** Log2FCs of PDH subunits for *h-bou* as compared to Col-0 enzymes. (A) Log2FC between *h-bou* and Col-0 for the lipoate containing E2 subunit (AT1G48030). (B) Log2FC between *h-bou* and Col-0 for subunit 2-1 (AT3G52200). (C) Log2FC between *h-bou* and Col-0 for the E1 subunit (AT1G59900); n = 3.
